# eIF4G2-Mediated Translation Initiation of Histone Modifiers Is Essential for Intestinal Stem Cell Maintenance and Differentiation

**DOI:** 10.1101/2025.05.19.652834

**Authors:** Haruko Kunitomi, Aye Myat Khaine, Radia Jamee, Vanessa Arreola, Mariselle Lancero, Amba Raychaudhuri, Samuel Perli, Yoshiko Sato, Mio Iwasaki, Pedro Ruivo, Kiichiro Tomoda, Mari Mito, Yuichi Shichino, Shintaro Iwasaki, Shinya Yamanaka

## Abstract

eIF4G2 (also known as NAT1, p97, or DAP5) is an evolutionally conserved protein homologous to the C-terminal two-thirds portion of eukaryotic translation initiation factor (eIF) 4G. Despite its abundant and ubiquitous expression, the physiological and pathological functions of eIF4G2 are poorly understood. Here, we show that acute loss of eIF4G2 in adult mice results in rapid weight loss with abnormalities in multiple organs, including impaired maintenance and differentiation of intestinal stem cells. Genome-wide ribosome profiling revealed that eIF4G2 is critical for the translation of key histone modification proteins involved in intestinal stemness. Our study underscores the importance of eIF4G2-mediated translation initiation in multicellular organisms.

## [Introduction]

Similar to transcription, the main regulatory checkpoint of protein translation occurs during initiation^1^. The generally accepted view of translation initiation is that a protein complex, which is orchestrated by the scaffold protein eukaryotic translation initiation factor (eIF) 4G, binds to the cap structure at the 5′-end of mRNA and scans 5′-untranslated regions (UTRs) toward the 3′-direction until it finds the methionine initiation codon^2^. The scaffold protein eIF4G was identified in 1982 as p220, and shown to be cleaved by picornaviral proteases into two fragments^3^. The cleaved N-terminal fragment binds to the cap-binding protein eIF4E, whereas the C-terminal fragment recruits ribosomes and other initiation factors, such as eIF3 proteins and eIF4A^4^. The cleavage, thus, results in the shutdown of cap-dependent translation of host mRNAs, whereas the cleaved C-terminal portion of eIF4G binds to specific sequences of viral RNAs, known as internal ribosomal entry site (IRES), to support cap-independent translation initiation^5^. Based on this historical finding, the vast majority of cellular mRNA is thought to be translated through eIF4G-mediated cap-dependent translation initiation.

In 1997, multiple groups identified a gene encoding a protein closely related to eIF4G. We identified the gene as NAT1 (Novel APOBEC1 Target #1), whose mRNA was aberrantly edited by overexpressed APOBEC1, an mRNA editing enzyme^6^. Three other groups designated the same gene as p97 based on the protein size^7^, or DAP5 (death-associated protein #5) based on its putative role in apoptosis^8^. Currently, the official nomenclature for the gene is eIF4G2, based on its homology to eIF4G^9^, whereas the original eIF4G is now designated eIF4G1. Importantly, eIF4G2 is homologous to only the C-terminal portion of eIF4G1, which is produced by the viral protease-mediated cleavage^9^. As predicted from its sequence, eIF4G2 binds to eIF3 proteins, eIF4A and ribosomal proteins, but not to the cap-binding protein eIF4E^6,7^.

Multiple roles for eIF4G2 in translation initiation have been proposed. Based on the similarity with cleaved eIF4G1, the most straightforward assumption is that eIF4G2 is involved in IRES-mediated translation initiation of cellular mRNAs^10–14^. Alternatively, eIF4G2 may recognize m^6^A-modified sequences adjacent to initiation codons^15–17^. Another model is that eIF4G2 is involved in cap-dependent translation through a partner protein other than eIF4E, namely eIF3d^18^. More recently, eIF4G2 has been proposed to engage in translation initiation of main open reading frames (ORFs) of mRNAs that contain upstream ORFs (uORFs) in their 5′ UTRs^19–21^. However, these studies relied on experiments performed *in vitro* or in cultured cells. The precise roles of eIF4G2 in translation initiation, especially *in vivo*, remain elusive.

The physiological and pathological roles of eIF4G2 also remain to be determined. We have generated knockout mice and showed that eIF4G2 is indispensable for early mouse development^22^. Without functional eIF4G2, mouse embryos cannot complete gastrulation. In addition, eIF4G2 is crucial for maintenance of pluripotency in mouse and human stem cells^23–25^. We also showed that loss of the *eIF4G2* homolog in *Drosophila* causes embryonic lethality and abnormal germband extension^26^. Because of its ubiquitous and abundant expression, as well as its evolutionary conservation from fly to human^27^, we predict that eIF4G2 has widespread and important roles in multicellular organisms. However, discovery of these roles has been stymied by the embryonic lethality of the knockout. Therefore, to precisely examine the functions of eIF4G2 and its roles in translation initiation *in vivo*, it was imperative to generate inducible gene knockout (KO) mouse models.

Here, we report Cre-LoxP-mediated conditional *Eif4g2*-KO mice. Our initial analysis of adult mice with acute loss of eIF4G2 demonstrated progressive weight loss and phenotypes in the small intestine, with loss of intestinal stem cells (ISCs). Through analysis of small intestinal organoids (SIOs) and the translatome, we revealed the critical roles of eIF4G2 in the small intestine. Our data *in vivo* and in SIOs collectively show that eIF4G2-dependent translation initiation of histone modifiers, such as CREBBP and EP300, is crucial for stemness in the small intestine. This study has revealed important roles of a general translation initiation factor in cell fate determination.

## [Results]

### Generation of a tamoxifen-inducible *Eif4g2*-KO mouse line

The mouse *Eif4g2* gene, located on chromosome 7, contains 22 exons spanning approximately 15 kilo base pairs (bps). To create a conditional *Eif4g2*-KO mouse line, we targeted the 103 bp-long exon 5, the deletion of which would cause a frameshift and premature stop codon. We inserted two loxP sites into the introns flanking exon 5 of the mouse *Eif4g2* gene by microinjecting CRIPSR/Cas9 ribonucleoproteins and single-stranded oligonucleotides to a C57BL/6 blastocyst (Figure 1A). After confirming successful generation of the floxed allele, we intercrossed the *Eif4g2*^flox/+^ mice and obtained *Eif4g2*^+/+^, *Eif4g2*^flox/+^, and *Eif4g2*^flox/flox^ offspring at predicted Mendelian ratios (Figure S1A), indicating that the gene editing did not affect viability or fertility. We then crossed the *Eif4g2*-floxed mouse line with B6.129-*Gt (ROSA)26Sor^tm^*^1^(cre/ERT2)*^Tyj^*/J (*Rosa26*-CreER) mice^28^ to create a tamoxifen-inducible systemic *Eif4g2*-KO mouse model. We treated the offspring with tamoxifen and extracted genomic DNA from various organs at day 14 post-treatment. With the exception of the brain, the PCR results showed complete gene KO in all investigated organs of tamoxifen-treated *Eif4g2*^flox/flox^, *Rosa26*-CreER (hereafter *Eif4g2*^-/-^) mice (Figure S1B). In intestinal tissue, protein analyses revealed complete loss of eIF4G2 protein as early as 6 days after tamoxifen treatment (Figure S1C), validating the generation of tamoxifen-inducible *Eif4g2*-KO mice.

**Figure 1.**
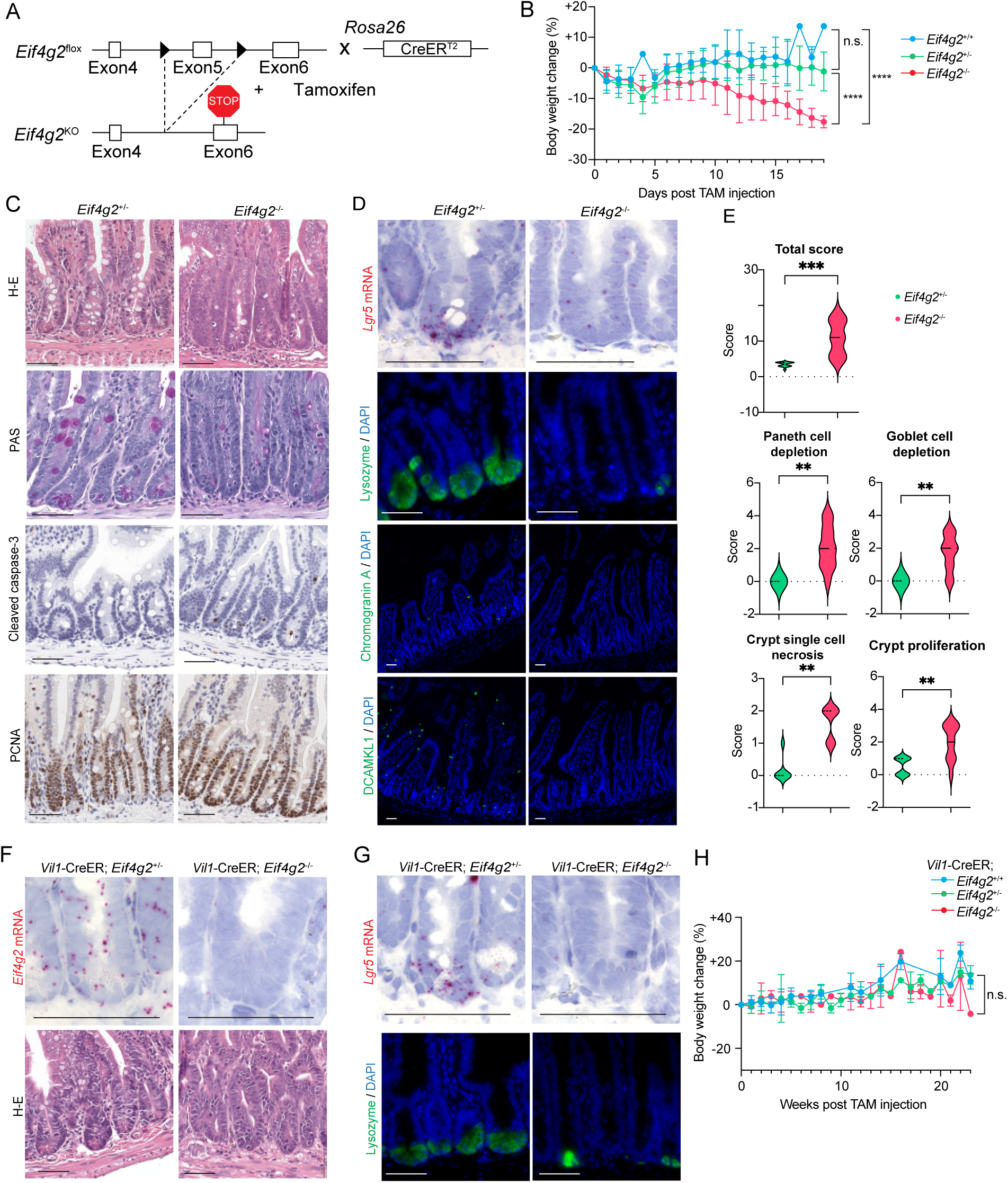
eIF4G2 is indispensable for ISCs in mice. (A) Schematic of the Tamoxifen-inducible mouse *Eif4g2* gene KO system. Black triangles represent loxP sites flanking the 103 bp long exon 5. (B) Body weight change after tamoxifen treatment in *Eif4g2*-floxed, *Rosa26*-CreER mice (wild type n=11, heterozygote n=24, homozygote n=29). (C) Histological findings in *Eif4g2*-floxed, *Rosa26*-CreER mice at endpoint. From top; Hematoxylin-Eosin (H-E), periodic acid-Schiff (PAS), cleaved caspase-3 and PCNA immunohistochemistry (IHC). (D) Histological analysis of *Eif4g2*-floxed, *Rosa26*-CreER mice at endpoint. (From top) RNAscope of the ISC marker *Lgr5*, fluorescent IHC of Paneth cell marker lysozyme, enteroendocrine cell marker chromogranin A and tuft cell marker DCAMKL1. (E) Violin plots showing modified intestinal lesion score of *Eif4g2*^+/-^ and *Eif4g2*^-/-^ (*Rosa26*-CreER) small intestines at end point (heterozygote n=13, homozygote n=11. See also Table S1). (F) Histological findings in *Vil1*-CreER; *Eif4g2*^+/-^and *Eif4g2*^-/-^ mice 1 month after tamoxifen treatment. Top; *Eif4g2* mRNA BaseScope, Bottom; H-E. (G) Histological findings in *Vil1*-CreER; *Eif4g2*^+/-^ and *Eif4g2*^-/-^ mice 1 month after tamoxifen treatment. Top; ISC marker *Lgr5* RNAscope, Bottom; Paneth cell marker lysozyme IHC. (H) Body weight change after tamoxifen treatment in *Eif4g2*-floxed, *Vil1*-CreER mice (wild type n=7, heterozygote n=7, homozygote n=5). * p <0.05, ** p < 0.005, *** p < 0.001, **** p < 0.0001. Scale bars indicate 50 um.

### eIF4G2 is indispensable for ISCs in mice

To investigate the effect of systemic *Eif4g2*-KO in adult mice, we treated 20-to 40-week-old mice with tamoxifen and observed their general condition and body weight over time. While *Eif4g2* wild type (*Eif4g2*^+/+^) and heterozygote-KO (*Eif4g2*^+/-^) mice showed no overt phenotypes, homozygote-KO (*Eif4g2*^-/-^) mice progressively lost weight and mobility and were euthanized at a humane endpoint of 15% weight loss within a few weeks (Figure 1B). At this early point, we evaluated the organ weight and histological findings throughout the body. *Eif4g2*^+/-^ mice did not show obvious pathological findings in any of the organs, in line with our previous study demonstrating normal development in conventional heterozygous KO mice^22^. In sharp contrast, *Eif4g2*^-/-^ mice showed significant abnormalities in multiple organs, such as bone marrow (Figure S1D) and small intestine. The current study focused on the latter since it could explain the rapid weight loss.

Hematoxylin-eosin (H-E) staining of the small intestine evidenced a decreased villus/crypt ratio in the *Eif4g2*^-/-^ mice, which is indicative of underlying damage, with associated scarce goblet and Paneth cells (Figure 1C). Periodic acid-Schiff (PAS) staining confirmed the marked decrease of goblet cells. In *Eif4g2*^-/-^ crypts, RNAscope in situ hybridization showed minimal *Lgr5* mRNA expression, and immuno-histochemistry barely detected lysozyme-positive Paneth cells, chromogranin A-positive enteroendocrine cells, or DCAMKL1-positive tuft cells (Figure 1D), indicating depletion of ISCs and all secretory cells. We detected increased numbers of apoptotic cells that were positive for cleaved caspase-3 in *Eif4g2*^-/-^ crypts (Figure 1C). Protein expression analyses of isolated intestinal crypts confirmed significant depletion of eIF4G2, OLFM4, and lysozyme (Figure S1C). In contrast to the depletion of ISCs, we observed a normal number of proliferating cell nuclear antigen (PCNA)-positive proliferating cells (Figure 1C). Together with the apparently normal villous structure, proliferation of the TA cells and their differentiation into enterocytes seemed to be maintained even in the absence of eIF4G2.

To more objectively quantify the histological findings, we blindly evaluated the tissue using a modified scoring system originally proposed by Erben et al^29^. In this system, each lesion was scored from 0 (normal) to 4 (most severe). The analyses confirmed that intestinal lesions were statistically significant in *Eif4g2*^-/-^ mice (Figure 1E and Table S1), including depletion of Paneth and goblet cells, accelerated necrosis in crypts, and increased crypt proliferation. Significant goblet cell loss was also observed in the colon (Figure S1E). In contrast to *Eif4g2*^-/-^ mice, *Eif4g2*^+/-^ mice did not show significant pathological findings (Figure 1C-1E). These findings demonstrated that eIF4G2 is indispensable for ISC maintenance and their differentiation towards the secretory lineage.

### Intestinal epithelium-specific *Eif4g2*-KO results in ISC depletion, but not to weight loss

The apparently normal epithelium in *Eif4g2*^-/-^ mice, despite the depletion of ISCs and secretory cells, was puzzling. To address the effect of eIF4G2 depletion for a prolonged period, we crossed the *Eif4g2*-floxed mice with B6.Cg-Tg(*Vil1*-Cre/ERT2)23Syr/J mice^30^ to generate an intestinal epithelium-specific *Eif4g2*-KO model. After tamoxifen treatment at 20 – 40 week age, RNA in situ hybridization by BaseScope using probes to detect *Eif4g2* mRNA confirmed successful gene ablation specifically in the intestinal epithelia (Figure 1F), with only a small fraction of crypts escaping gene KO at one-month post-injection (Figure S1F). The intestinal phenotype seen in *Eif4g2*^-/-^ mice was fully recapitulated in intestinal epithelium-specific KO mice; intestinal epithelium and proliferation of TA cells were maintained despite severe depletion of ISCs and all secretory lineage cells, and increased apoptosis was observed in the crypt (Figure 1G, Figure S1G-I, Table S2). The intestinal epithelium-specific *Eif4g2*^-/-^ mice did not lose weight (Figure 1H), suggesting that the weight loss observed in the global KO was not attributable to the intestinal abnormality.

### The *Eif4g2*-KO epithelium exhibits fetal-like reversion

We next tried to understand how enterocytes were maintained in the absence of ISCs in the eIF4G2-depleted intestine. To this end, we performed RNA-seq on intestinal crypt and villous tissues. In line with the histological findings, ISC marker genes and all secretory lineage marker genes were depleted in the *Rosa26-CreER; Eif4g2*^-/-^ crypts (Figure 2A). Expression of homeobox transcription factors (*Cdx1*, *Cdx2*)^31^, and pioneer transcription factors (*Foxa2*, *Foxa3*)^32^, all of which contribute to intestinal lineage specification, was significantly reduced in the *Eif4g2*^-/-^ crypts. We observed upregulation of markers for non-intestinal lineages, such as mesenchymal (*Foxc1*, *Vgll3*, *Nanos1*), immune (*Gbp2*, *Gbp3*, *Slfn5*), and endothelial (*Cav1*, *Cavin2*, *Cav2*) (Figure 2A). In sharp contrast, the *Eif4g2*^-/-^ villi exhibited increased expression of immature and mature enterocyte markers (Figure 2B), confirming the paradoxical preservation of enterocytes despite the loss of ISCs.

**Figure 2:**
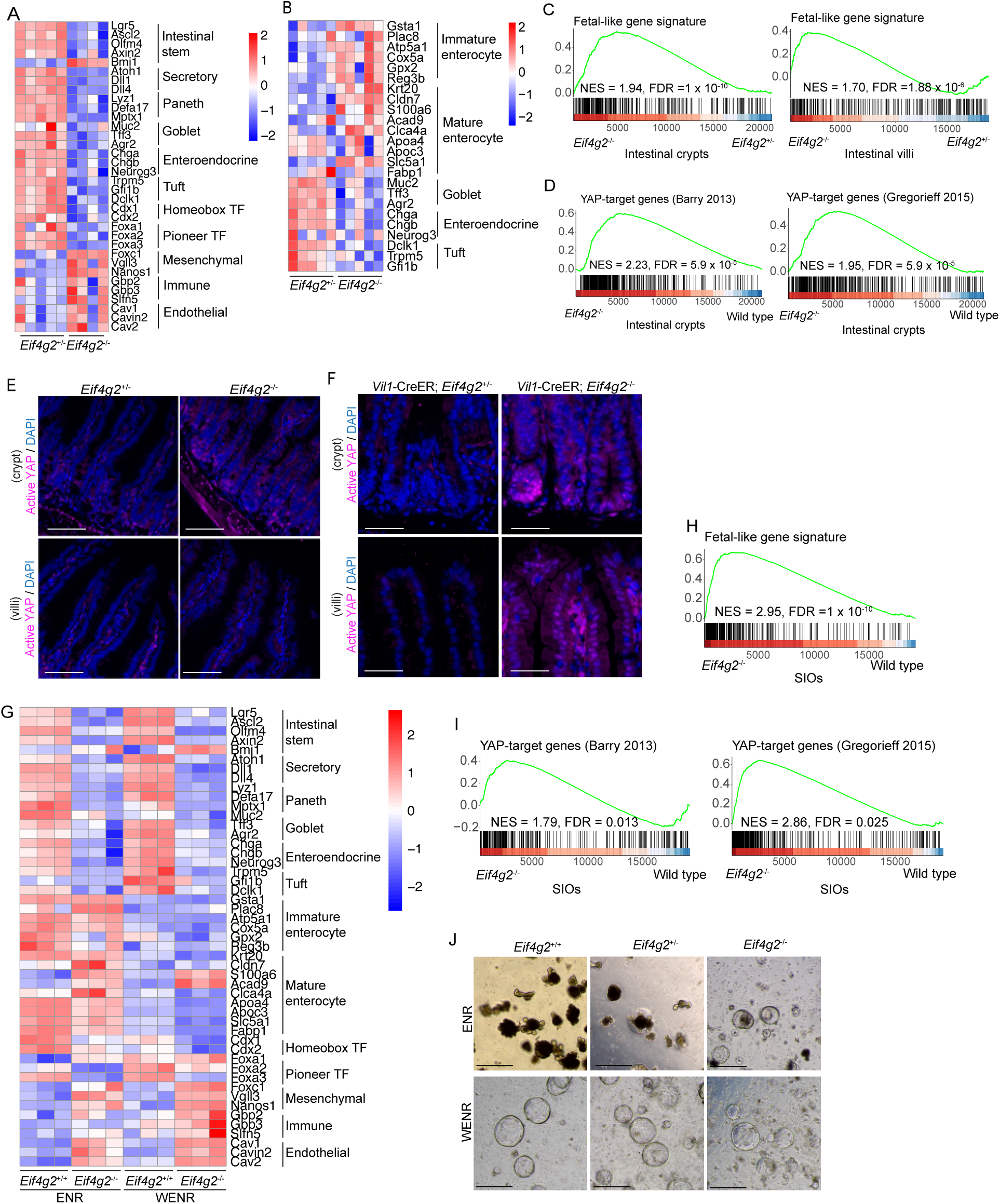
*Eif4g2*-KO intestines exhibit fetal-like reversion. (A) Heatmap showing representative gene expression of each cell type in *Eif4g2*^+/-^ and *Eif4g2*^-/-^ intestinal crypts. (B) Heatmap showing representative gene expression of each cell type in *Eif4g2*^+/-^ and *Eif4g2*^-/-^ intestinal villi. (C) GSEA showing enrichment of the fetal-like gene signature in *Eif4g2*^-/-^ (Left) crypts and (Right) villi. (D) GSEA showing enrichment of YAP-target gene signature in *Eif4g2*^-/-^ crypts. (E) Active YAP IHC in *Rosa26*-CreER; *Eif4g2*^+/-^ and *Eif4g2*^-/-^ mice 14 days after tamoxifen treatment. Top; crypts, bottom; villi. (F) Active YAP IHC in *Vil1*-CreER; *Eif4g2*^+/-^ and *Eif4g2*^-/-^ mice 1 month after tamoxifen treatment. Top; crypts, bottom; villi. (G) Heatmap showing representative gene expression of each cell type in *Eif4g2*^+/+^ and *Eif4g2*^-/-^ SIOs. Left half shows data from ENR condition and right shows WENR condition. (H) GSEA showing enrichment of the fetal-like gene signature in *Eif4g2*^-/-^ SIOs grown in ENR condition. (I) GSEA showing enrichment of YAP-target gene signature in *Eif4g2*^-/-^ SIOs grown in ENR condition. (J) Representative morphology of *Eif4g2*^+/+^, *Eif4g2*^+/-^ and *Eif4g2*^-/-^ SIOs grown in (top) ENR condition and (bottom) WENR condition 20 days after 4-OHT treatment. In (E) and (F), scale bars indicate 100 um. In (J), scale bars indicate 500 um.

Loss of ISCs after radiation, diphtheria toxin-mediated ISC ablation, and chemotherapy is known to trigger a transient “fetal reversion”, in which multiple progenitors and differentiated cell types dedifferentiate to rebuild the stem-cell pool^33–35^. Accordingly, *Eif4g2*^-/-^ crypts exhibited a pronounced enrichment of the ‘fetal-like crypt gene signature’^36^ (NES = 1.94, FDR <0.01, Figure 2C, left). Within this paradigm, activation of YAP signaling is a key driver^35^. Consistent with this observation, our GSEA demonstrated significant enrichment of two independent YAP-target signatures^37,38^ in *Rosa26*-CreER; *Eif4g2*^-/-^ crypts 14 days after TAM treatment (Figure 2D), although nuclear YAP was not yet detectable by immunostaining at that time (Figure 2E) – a lag that has been reported in other regenerative contexts^33,38^. One month after TAM treatment, intestinal epithelium-specific *Eif4g2*^-/-^ crypts displayed pronounced nuclear YAP accumulation (Figure 2F), indicating that YAP activation persists and amplifies over time. These findings suggest that loss of eIF4G2 triggers an early transcriptional activation of YAP that subsequently manifests as robust nuclear YAP, thereby sustaining the prolonged fetal-like reversion of the crypts.

Consistent with the fetal-like reversion paradigm, we found that *Bmi1*—originally described as an adult +4 reserve ISC marker^39^ and more recently recognized as a hallmark of fetal ISCs^40^—, was markedly upregulated in the *Eif4g2*^-/-^ crypts (Figure 2A). RNAscope in situ hybridization revealed abundant *Bmi1* transcripts along the entire length of the *Eif4g2*^-/-^ crypt, whereas in *Eif4g2*^+/-^ controls was confined to the occasional quiescent *Bmi1+* cells (Figure S2A). Semi-quantitative scoring (0-4 scale) confirmed a significant genotype effect (mixed-effects two-way ANOVA, F = 79.0, P <0.0001) and higher scores in the KO at multiple positions after Sidák correction (Figure S2B), indicating that *Bmi1* is expressed both more strongly and across a broader positional domain in *Eif4g2*^-/-^ crypts. Given that the *Bmi1*^+^ ISCs in the embryonic day 12.5 fetal gut are fast-cycling and multipotent cells yet molecularly distinct from *Lgr5*^+^ ISCs, our data raise the possibility that loss of eIF4G2 either induces or expands a fetal-type *Bmi1*^+^ ISC population in the adult crypt.

The ‘fetal-like crypt gene signature’ was also enriched in the *Eif4g2*^-/-^ villi (NES = 1.70, FDR < 0.01, Figure 2C, right). Correspondingly, Active YAP staining was also present in villus epithelial nuclei of the intestinal epithelium-specific *Eif4g2*^-/-^ mice 1 month after TAM treatment (Figure 2F), a hallmark of atrophy-induced villus epithelial cells (aVEC)^41^.

Collectively, our findings suggest that eIF4G2 deficiency triggers an adaptive, fetal-like regeneration program across the entire epithelium: crypts are repopulated by fast-cycling *Bmi1*^+^ fetal-like ISCs, while villus enterocytes shift towards an aVEC-like state. This dual response compensates for the depletion of canonical *Lgr5*^+^ ISCs and helps maintain epithelial integrity.

### Small intestinal organoids recapitulate the in vivo findings

The crypt-villi structure and rapid turnover can be recapitulated in SIOs, grown in Matrigel-based culture with medium containing epithelial growth factor (EGF), Noggin, and Rspondin-1 (ENR)^42^. To further investigate the roles of eIF4G2 in intestine, we generated SIOs from *Rosa26*-CreER; *Eif4g2*^+/+^, *Eif4g2*^flox/+^, and *Eif4g2*^flox/flox^ mice. We confirmed the complete depletion of eIF4G2 protein within 3 days after 4-hydroxytamoxifen (4-OHT) treatment in SIOs from *Rosa26*-CreER; *Eif4g2*^flox/flox^ mice (Figure S2C). RNA-seq analyses at 6 days after 4-OHT revealed downregulation of ISC and secretory cells markers in *Eif4g2*^-/-^ SIOs, while enterocyte markers remained largely unchanged (Figure 2G, left). Consistent with the findings *in vivo* (Figure 2A), we observed a similar transcriptional shift in *Eif4g2*^-/-^ SIOs, including reduced expression of homeobox transcription factors (*Cdx1*, *Cdx2*) and pioneer transcription factors (*Foxa2, Foxa3*), accompanied by increased expression of mesenchymal (*Foxc1*, *Vgll3*, *Nanos1*), immune (*Gbp2*, *Gbp3*, *Slfn5*), and endothelial (*Cav1*, *Cavin2*, *Cav2*) related genes (Figure 2G, left). These changes suggest a loss of the intestinal epithelial gene signature in the absence of eIF4G2. Furthermore, an enrichment of the ‘fetal-like crypt gene signature’ was also observed (NES = 2.96, FDR <0.01, Figure 2H). GSEA confirmed YAP-target enrichment (Figure 2I), and immunoblotting demonstrated elevated active YAP (Figure S2D), including cystic regions normally devoid of nuclear YAP (Figure S2E). *Eif4g2*^-/-^ SIOs gradually lost the budding structure and became cystic within 20 days after 4-OHT treatment (Figure 2J, top), but were still expandable up to 30 days. This is in sharp contrast to Lgr5-depleted SIOs that can propagate only for a several days^43^, further confirming the prolonged survival of eIF4G2-depeleted enterocytes.

To further investigate the role of eIF4G2 in ISC differentiation, we treated SIOs with small molecules that modulate self-renewal and lineage specification^44^. Briefly, ISCs were first expanded in ENR medium supplemented with glycogen synthase kinase 3 (GSK3) inhibitor CHIR99021 and Valproic acid, after which 4-OHT was added to knockout eIF4G2 and lineage-directed differentiation was induced by combinational activation or inhibition of Wnt and Notch signaling (CHIR99021/DAPT, IWP-2/DAPT, or IWP-2/Valproic acid). In *Eif4g2*^+/+^ SIOs, these regimens efficiently upregulated marker genes for Paneth cells (*Lyz1*, *Mptx1*), goblet cells (*Muc2*, *Clca1*), and enterocytes (*Alpi*, *Fabp1*) (Figure S2F). By contrast, Paneth cell and goblet cell differentiation was significantly impaired in *Eif4g2*^-/-^ SIOs, whereas enterocyte differentiation was largely preserved (Figure S2G). Notably, RNA-seq of both *Eif4g2*^-/-^ crypts and SIOs revealed a profound reduction in the transcript level of *Atoh1* (Figure 2A and 2G), the master regulator of secretory-lineage commitment^45^. The inability of potent chemical cues to rescue secretory differentiation therefore underscores an essential requirement for eIF4G2 in secretory-lineage specification.

### Wnt activation rescues ISC survival but does not restore lineage commitment and differentiation

Since Paneth cells function as the stem cell niche that provides critical signals such as Wnt ligands^46^, we next investigated whether their loss was responsible for ISC depletion in the *Eif4g2*^-/-^ intestine. To this end, we cultured SIOs in the Wnt3a-supplemented (WENR) condition that is capable of supporting stem cell survival without Paneth cells^47^. Wnt3a addition enabled long-term survival of *Eif4g2*^-/-^ SIOs for over 10 passages, with proliferation and cystic morphology comparable to *Eif4g2*^+/+^ and *Eif4g2*^+/-^ SIOs (Figure 2J bottom), suggesting that Wnt signaling activation prolonged the survival of *Eif4g2*^-/-^ ISCs. However, RNA-seq demonstrated that even under WENR condition, *Eif4g2*^-/-^ SIOs failed to restore normal gene expression of ISC and secretory cell markers (Figure 2G, right). In addition, expression of homeobox transcription factors (*Cdx1*, *Cdx2*), pioneer transcription factors (*Foxa2*, *Foxa3*), mesenchymal (*Foxc1*, *Vgll3*, *Nanos1*), immune (*Gbp2*, *Gbp3*, *Slfn5*), and endothelial (*Cav1*, *Cavin2*, *Cav2*) related genes showed similar patterns to *Eif4g2*^-/-^ crypts and organoids in ENR condition. These data demonstrate that Wnt signaling activation alone is insufficient to restore intestinal lineage identity in the *Eif4g2*^-/-^ organoids.

### Translation efficiency of epigenetic regulators is suppressed by eIF4G2 depletion

To investigate the molecular basis of ISC signature loss, we performed paired Ribo-seq^48^ and RNA-seq in *Eif4g2*^+/+^, *Eif4g2*^+/-^, and *Eif4g2*^-/-^ SIOs in the ENR condition at day 4 after 4-OHT treatment. Translation efficiency (TE) was estimated by normalizing Ribo-seq read counts with RNA-seq levels. As expected, *Eif4g2*^+/-^ SIOs showed minimal differences in TE compared to *Eif4g2*^+/+^ (Figure 3A). In sharp contrast, *Eif4g2*^-/-^ SIOs exhibited significant TE decrease in 283 genes and increase in 155 genes (Figure 3B, Table S3).

**Figure 3:**
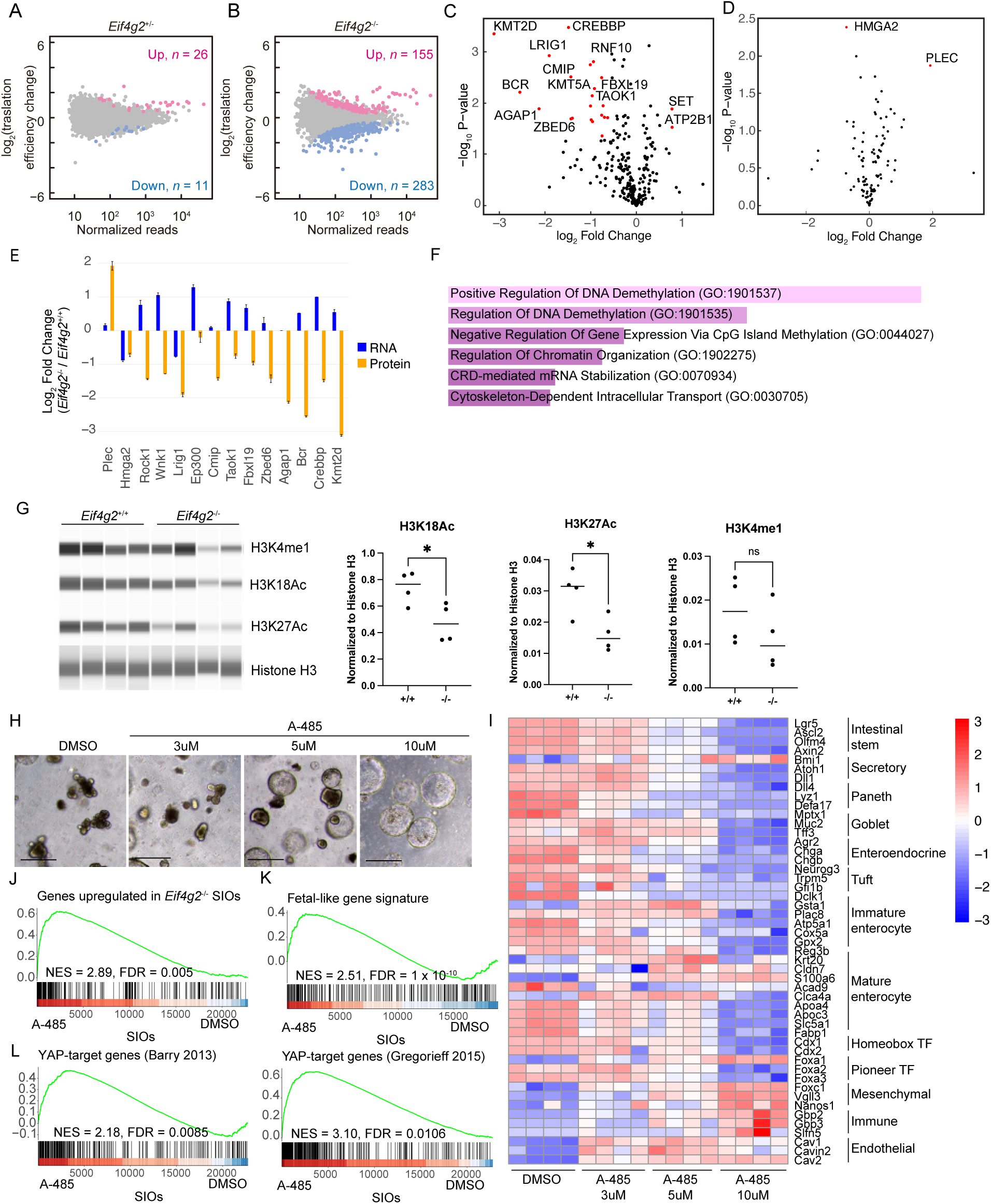
*Eif4g2*-KO causes epigenetic dysregulation and leads to ISC loss. (A, B) MA plot showing translation efficiency (TE) changes between (A) *Eif4g2*^+/-^ and *Eif4g2*^+/+^ and (B) *Eif4g2*^-/-^ and *Eif4g2*^+/+^ SIOs. Red and blue dots indicate genes with significantly higher and lower TE, respectively (false discovery rate < 0.05). (C, D) Volcano plots of protein expression changes in *Eif4g2*^-/-^ and *Eif4g2*^+/+^ SIOs 6 days after 4-OHT treatment showing genes with significantly lower (C) and higher (D) TE upon *Eif4g2*-KO. (E) Comparison of RNA and protein expression change in selected ‘eIF4G2-dependent’ genes, based on RNA-seq and proteomics data. (F) Gene ontology analysis results of ‘eIF4G2-dependent’ genes. (G) Protein quantification of H3K4me1, H3K18ac, H3K27ac, and Histone H3 in *Eif4g2*^+/+^ and *Eif4g2*^-/-^ SIOs 6 days after 4-OHT treatment. n=4 each. (Left) Wes run image, (Right) quantification of proteins normalized to Histone H3 expression. Bars indicate the mean of the replicates. (H) Representative morphology of SIOs treated with DMSO or A-485 (3, 5, 10uM) for 4 days. (I) Heatmap showing representative gene expression of SIOs treated with DMSO or A-485 (3, 5, 10uM) for 4 days. (J) GSEA showing enrichment of genes upregulated upon *Eif4g2*-KO in SIOs treated with 10uM A-485 for 4 days. (K) GSEA showing enrichment of the fetal-like gene signature in SIOs treated with 10uM A-485 for 4 days. (L) GSEA showing enrichment of YAP target genes in SIOs treated with 10uM A-485 for 4 days. * p <0.05. Scale bars indicate 500um.

To assess whether changes in TE were reflected at the protein level, we performed proteomic analyses on day 6 after 4-OHT treatment of *Eif4g2*^+/+^ and *Eif4g2*^-/-^ SIOs. We used the WENR condition to obtain sufficient materials in the downstream study. We compared the protein expression changes among genes with significantly reduced TE, increased TE, and unchanged TE between *Eif4g2*^-/-^ and *Eif4g2*^+/+^ SIOs. We found that genes with decreased TE exhibited significantly lower protein abundance even at this early time point (Figure 3C), whereas those with increased TE did not show corresponding increases in protein levels (Figure 3D). We focused on the genes with significantly decreased TE and designated them ‘eIF4G2-dependent’ genes (Table S3). Comparison with paired RNA-seq data confirmed that the decrease of these proteins, including epigenetic gene regulators CREBBP (cyclic AMP response element-binding protein binding protein), EP300, and KMT2D, was at the post-transcriptional level (Figure 3E). The eIF4G2 dependencies in a subset of genes, such as *Eif4g2*, *Crebbp*, *Kmt2d*, *Rock1*, *Wnk1*, and *Ep300*, have previously been reported in mouse and human pluripotent stem cells^23,49^, suggesting a common role for eIF4G2 in the translation initiation of these genes in different cell types and species. Western blot analyses with capillary electrophoresis confirmed a decrease in CREBBP and EP300 proteins levels at day 6 after 4-OHT treatment in *Eif4g2*^-/-^ SIOs (Figure S3A). Gene ontology analyses of the ‘eIF4G2-dependent’ genes showed enrichment of genes related to DNA demethylation, chromatin regulation, mRNA stabilization, and cytoskeleton-dependent intracellular transport (Figure 3F). These results prompted us to further investigate epigenetic status in *Eif4g2*^-/-^ SIOs.

### Intestinal cells lose H3K18 and H3K27 acetylation upon *eIF4G2* depletion

CREBBP and EP300 are the two members of the KAT3 subfamily of histone acetyltransferases (HATs) with preferences towards H3K18 and H3K27 acetylation^50^. KMT2D is the major mammalian H3K4 mono- and di-methyltransferase that marks active enhancers during differentiation^51^. To evaluate the effects of the decreased protein levels of CREBBP, EP300, and KMT2D, we quantified the abundance of H3K18ac, H3K27ac, and H3K4me1 in *Eif4g2*^+/+^ and *Eif4g2*^-/-^ SIOs at 6 days (Figure 3G) and 18 days (Figure S3B) after 4-OHT treatment. Both H3K18 and H3K27 acetylation were significantly reduced in *Eif4g2*^-/-^ SIOs by day 6 and further decreased by day 18, indicating the progressive loss of histone acetylation on these lysine residues after eIF4G2 depletion. H3K4me1 was also decreased by day 18 after *Eif4g2*-KO, but the change was slower and not significant at day 6. Since ISC loss occurs as early as day 6 in *Eif4g2*^-/-^ SIOs, it may be attributable to the decreased protein levels of CREBBP and EP300.

### CREBBP/EP300 inhibitors recapitulate the eIF4G2 knockout phenotype

To investigate whether the decrease in CREBBP/EP300 underlies the loss of ISCs in *Eif4g2*^-/-^ SIOs, we treated wild-type SIOs with the small molecule A-485, which specifically inhibits CREBBP/EP300 through competition at the acetyl-CoA binding site^52^. A-485 treatment dose-dependently reduced the acetylation marks on H3K18 and H3K27 residues in SIOs (Figure S3C) and induced cystic morphological changes similar to those seen in *Eif4g2*^-/-^ SIOs (Figure 3H). Furthermore, key gene expression patterns in A-485 treated SIOs matched those of *Eif4g2*^-/-^ SIOs, with decreased expression of all intestinal epithelial cell type markers, downregulation of homeobox transcription factors and pioneer transcription factors, and upregulation of mesenchymal, immune, and endothelial cell markers in a dose-dependent manner (Figure 3I, S3D). GSEA revealed that A-485 caused strong enrichment of genes upregulated by loss of eIF4G2 (NES = 2.89, FDR < 0.01, Figure 3J). A-485 treatment also recapitulated the enrichment of the ‘fetal-like crypt gene signature’ (NES = 2.51, FDR < 0.01, Figure 3K) and YAP-target genes (Figure 3L), and induced marked YAP signaling activation (Figure S3E, Figure S3F). These results indicate that the decreased protein levels of CREBBP and EP300 contribute, at least in part, to the ISC loss in the absence of eIF4G2.

### Ribosomal complexes are sequestered in long 5′ UTRs of ‘eIF4G2-dependent’ transcripts upon eIF4G2 loss

We investigated the characteristics of ‘eIF4G2-dependent’ genes and found that they had significantly longer 5′ UTR length than did the other genes (Figure 4A). Naturally, these ‘eIF4G2-dependent’ transcripts also had a significantly higher probability of having uORFs (Figure 4B). Interestingly, the ratio of ribosome-protected footprints (RPFs) on the 5′ UTR and those on the coding sequence (5′ UTR/CDS ratio) of ‘eIF4G2-dependent’ genes was significantly higher in *Eif4g2*^-/-^ SIOs (Figure 4C). This trend was clearly exemplified in *Crebbp* and *Ep300*; in *Eif4g2*^-/-^ SIOs, RPFs decreased on the CDS of these genes, whereas RPFs on their 5′ UTRs increased significantly (Figure 4D). A closer look at RPFs suggested that near cognate initiation codons may be translated in 5′ UTR of *Crebbp* and *Ep300* (Figure 4E) in *Eif4g2*^-/-^ SIOs. Collectively, these findings suggest that the 5′ UTR context is the key determinant for eIF4G2-dependent translation regulation and that eIF4G2 may be involved in leaky scanning or reinitiation after translation of uORFs.

**Figure 4:**
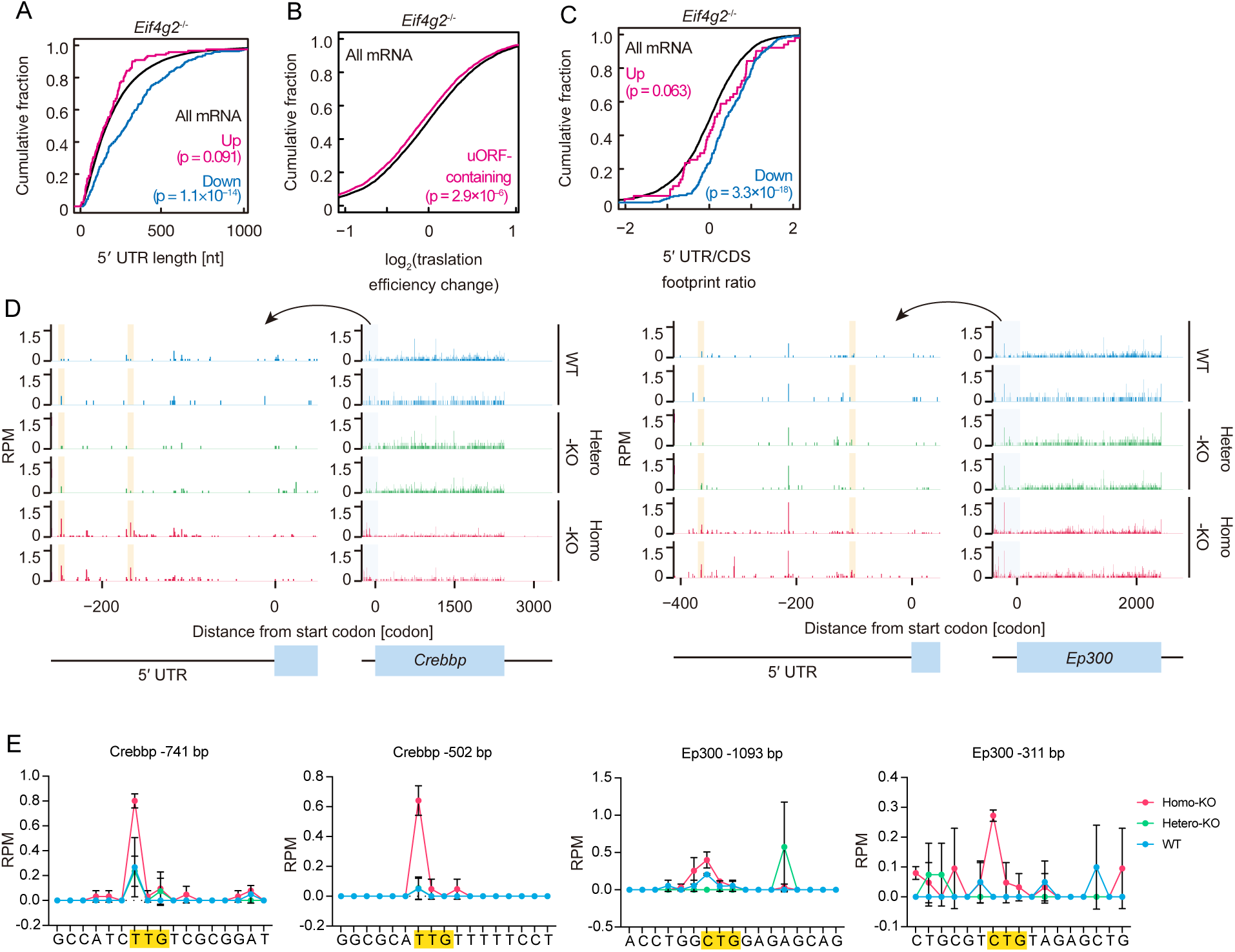
Ribosomal complexes are sequestered in long 5′ UTRs of eIF4G2-dependent genes upon eIF4G2 loss. (A) Comparison of 5′ UTR length of differentially translated genes in *Eif4g2*^-/-^ vs. *Eif4g2*^+/+^. Red and blue lines indicate genes with significantly higher and lower TE in the KO condition. (B) Frequency of uORFs in differentially translated genes in the *Eif4g2*^-/-^ vs. *Eif4g2*^+/+^ comparison. (C) 5’UTR/CDS ratio of RPFs mapping on ‘eIF4G2-dependent’ genes with significantly higher (red) and lower (blue) TE in *Eif4g2*^-/-^ SIOs. (D) Distribution of RPFs in the mouse (top) *Crebbp* (XM_006521751.5) and (bottom) *Ep300* (NM_177821.7) 5′ UTR and CDS. The number of RPFs with P-sites located on each nucleotide is plotted. Regions shown in (E) are highlighted. (E) Distribution of RPFs along the indicated sites in 5′ UTR. The number of RPFs with P-sites located on each nucleotide is plotted. Potential near-cognate start codons are highlighted in yellow. RPM, reads per mapped million reads. For (A)-(C), p values are calculated by the Mann-Whitney U test (two-tailed).

## [Discussion]

In this study, we generated a tamoxifen-inducible knockout model to examine the physiological roles of eIF4G2 in adult mice and to identify its in vivo target mRNAs. By leveraging this mouse model alongside intestinal organoid culture techniques, we identified eIF4G2 as a pivotal regulator of intestinal stem cell maintenance and differentiation. We uncovered a set of mRNAs whose translation depends on eIF4G2. Notably, eIF4G2 loss led to decreased translation of epigenetic modifiers and altered histone acetylation, underscoring eIF4G2’s critical role in maintaining intestinal homeostasis. These findings provide key insights into translational control in adult stem cells and open avenues for exploring the broader implications of eIF4G2 in tissue regeneration.

Upon eIF4G2 deletion, protein levels of CREBBP and EP300 were significantly reduced, though not abolished. Their mRNAs possess long 5′ UTRs with potential uORFs. Ribosome profiling in SIOs suggested that eIF4G2 facilitates leaky scanning or reinitiation after translation of these uORFs. In the absence of eIF4G2, translation initiation of the main ORFs was substantially impaired, though initiation via the eIF4G1-containing scanning complex still occurred at reduced efficiency. Notably, double heterozygosity for *Crebbp* and *Ep300* invariably results in embryonic lethality, with phenotypes resembling those of *Crebbp* or *Ep300* homozygous mutants^53,54^—highlighting the sensitivity of mouse development to the combined dosage of these closely related lysine acetyltransferases (KATs). Even single heterozygosity results in partial embryonic lethality, indicating that even a ∼25% reduction in CREBBP or EP300 can disrupt gene networks critical during development^53^. These observations argue that eIF4G2-dependent translation of *Crebbp* and *Ep300* is essential for normal cellular function mediated by these KATs.

Our results emphasize the significance of CREBBP/EP300 (also known as KAT3A and KAT3B) in ISCs. This is consistent with previous findings showing that mouse ISCs maintain an open chromatin state, characterized by H3K4me2, H3K27ac, and DNase I hypersensitivity^55^. While the roles of CREBBP/EP300 in ISCs were not previously characterized, KAT2A and KAT2B— which catalyze acetylation at H3K9 and H3K14—are known to support ISC maintenance^56^. Furthermore, valproic acid, a known HDAC1/2 inhibitor and Notch activator, promotes ISC proliferation in SIOs^44^. However, double knockout of HDAC1/2 results in ISC loss^57^, indicating that a proper level of histone acetylation is crucial for ISC function.

The ISC and secretory cell loss, along with fetal-like reversion observed in *Eif4g2* KO mice, was recapitulated by treatment with the CREBBP/EP300 KAT domain inhibitor A-485, supporting a functional link between eIF4G2-mediated translation and histone acetylation. In contrast, other inhibitors such as SGC-CBP30 and I-CBP112—targeting the bromodomain of CREBBP/EP300—were reported to enrich ISCs and enhance SIO growth^58^. A-485 mimics CREBBP/EP300 gene knockout in global acetylome profiles, whereas I-CBP112 showed much smaller effects^59^. Interestingly, I-CBP112 may even enhance CREBBP/EP300 activity^60^. These results may suggest that the KAT activity of CREBBP/EP300 is crucial for normal ISCs.

CREBBP/EP300 have broad functions beyond histone acetylation, including acetylation of proteins such as p53^61^ and STAT3^62^. Acetylome analyses following gene knockout or A-485 treatment revealed that CREBBP/EP300 acetylate thousands of sites across diverse signaling and regulatory proteins, including Notch, Wnt, and Hippo pathways^59^. In contrast, bromodomain inhibitors like I-CBP112 affected only a small subset of these sites. Our findings that A-485 phenocopied the intestinal defects of *Eif4g2*-KO, while SGC-CBP30 and I-CBP112 enhanced ISC maintenance^58^, suggest that loss of non-histone protein acetylation may also contribute to ISC depletion.

The *Eif4g2*-deleted intestine displayed a unique phenotype: despite near-complete loss of *Lgr5*⁺ ISCs and secretory cells, enterocytes remained intact. This was especially evident in intestine-specific KO mice, which retained normal body weight and morphologically intact villi for months despite lacking *Lgr5*⁺ ISCs. In contrast, diphtheria toxin-mediated ISC ablation caused rapid crypt loss and lethality^43^. In that study, SIOs failed to expand. Recent studies suggest that a fetal-like reversion mechanism enables recovery after transient *Lgr5*⁺ ISC ablation (reviewed by Viragova & Klein^35^), where fetal-like stem cells temporarily supply both enterocytes and secretory cells. In *Eif4g2*-depleted intestines and SIOs, we observed activation of a fetal-like gene program. These cells may compensate for the loss of *Lgr5*⁺ ISCs by continuously producing enterocytes. Furthermore, *Atoh1* expression was suppressed, which may explain the absence of secretory cells, as *Atoh1* depletion is known to convert secretory cells into functional enterocytes^55^.

The fetal-like reversion may be driven by YAP/TAZ and SOX9 transcription factors^35^. In line with this, both ‘fetal-like gene signature’ and YAP-target genes were significantly enriched in intestinal crypts following loss of eIF4G2. CREBBP/EP300 have been reported to enhance YAP activity via acetylation and direct interaction with the YAP/TEAD complex^63,64^. Notably, A-485 treatment activated YAP and induced fetal-like reversion, raising questions about whether CREBBP/EP300 act upstream, downstream, or in parallel with YAP in regulating intestinal regeneration and homeostasis.

We also observed increased *Bmi1*⁺ cells in the crypts of *Eif4g2*-KO intestines. Initially identified as +4 label-retaining cells, *Bmi*1-expressing cells are considered a distinct ISC population^39,65^. More recently, *Bmi1*⁺ cells were proposed to be predominant during early development, giving rise to *Lgr5*⁺ ISCs as development proceeds^40^. The emergence of *Bmi1*⁺ cells in the KO intestine may reflect an alternative ISC population arising through fetal-like reversion. The *Eif4g2*-KO model offers a valuable system to study intestinal cell plasticity.

## Limitations of study

While our data highlight CREBBP/EP300 as key mediators, additional eIF4G2-dependent targets may also contribute to the intestinal phenotype. We identified hundreds of such mRNAs, and their individual roles warrant further study. Moreover, the physiological functions of eIF4G2 in tissues beyond the intestine remain unclear. Mechanisms underlying the rapid weight loss observed in the systemic KO mice also remain to be elucidated. It is notable that intestinal epithelial and hematopoietic cells—two of the fastest-turnover cell types^66,67^—were affected within 2–3 weeks post-KO, at which point mice were euthanized due to weigh loss. Tissue- and organ-specific knockout models will be essential to clarify the broader roles of eIF4G2.

## Acknowledgements

We thank Dr. Toshiro Sato for scientific discussion. This work was supported by the Core Center for iPS Cell Research, Research Center Network for Realization of Regenerative Medicine, AMED under grant number JP21bm0104001 (S.Y.); iPS Cell Research Fund (S.Y.); KAKEHNHI under grant number JP24H02307 (S.I.); AMED under grant number JP20gm1410001 (S.I.) and JP23gm6910005 (Y.S.); and RIKEN Pioneering Projects (S.I. and Y.S.). This work was also supported by the on-site laboratory initiative launched by Kyoto University, funding from Mr. Hiroshi Mikitani, Mr. Marc Benioff, the L. K. Whittier Foundation, the Roddenberry Foundation, and Gladstone Institutes. We thank Gladstone Institutes Transgenic Core, Gladstone Institutes Microscopy and Histology Core, Gladstone Institutes Bioinformatics Core and the Comparative Pathology Laboratory (UC Davis) for technical assistance. This work uses the HOKUSAI SailingShip supercomputer facility at RIKEN.

## Author contributions

Conceptualization, H.K., S.P., K.T. and S.Y.; Methodology, H.K., M.I., P.R.R.,Y.S., and S.I.; Investigation, H.K, A.M.K., R.J., M.L., E.R., Y.S., M.I., P.R.R., K.T., M.M., Y.S., and S.I.; Data Curation, H.K., Y.S., and M.I.; Writing – Original Draft, H.K.; Writing – Review & Editing, H.K., M.I., Y.S., S.I., and S.Y.; Visualization, H.K. and Y.S.; Funding Acquisition, Y.S., S.I., and S.Y.; Supervision, S.Y.

## Supplemental information

Document S1. Figures S1-S3 and Tables S1-S3.

## [Methods]

### Animals

All protocols concerning animal use were approved by the Institutional Animal Care and Use Committees at the University of California, San Francisco and conducted in strict accordance with the National Institutes of Health Guide for the Care and Use of Laboratory Animals. All mouse strains were maintained on a predominantly C57BL/6 background. Animals were housed under specific pathogen-free, temperature- and humidity-controlled facility under a 12-hour light/dark cycle. Food and water were provided *ad libitum*. Mice were 20 to 40 weeks old at the time of knockout experiments, and at least 8 weeks old for cell isolations. Mice of both sexes were used in all experiments with littermate controls.

*Eif4g2^flox^* mice were produced by blastocyst injection of ribonucleoprotein (RNP) complexes consisting of purified Cas9 protein, 2 guide RNAs (crRNA targeting *Eif4g2* locus and universal 67-mer tracrRNA, and a single-stranded oligonucleotide DNA template for homology-directed repair (all from Integrated DNA Technologies (IDT) that led to the insertion of 2 loxP sites flanking the exon 5 of the *Eif4g2* gene. Blastocysts were transferred to pseudopregnant females and pups were weaned at 4 weeks of age. Founder animals were screened for introduction of the loxP sites by PCR amplification and confirmed by sequencing. Positive founders were outcrossed to wild-type C57BL/6J mice and pups were screened for germline transmission of the *Eif4g2^flox^* allele by PCR genotyping with sequencing confirmation. To generate tamoxifen-inducible models, we crossed the *Eif4g2^flox^* mice with *Rosa26-CreER* and *Vil1-CreER* mice. Mouse sources and citations are provided in the Materials Table.

To knock out floxed *Eif4g2* alleles in both *Rosa26*-CreER; *Eif4g2*^flox^ and *Vil1*-CreER; *Eif4g2*^flox^ mice, we administered 100mg/kg body weight tamoxifen (Sigma-Aldrich, stock solutions prepared in cornflower oil) by intraperitoneal (IP) injection daily for 4 consecutive days. Mice were weighed and observed daily.

### DNA extraction and genotyping with PCR

Mouse tissue was lysed in DirectPCR lysis reagent and proteinase K solution (both from Viagen Biotech) and digested overnight at 56°C. PCR reactions were carried out with KOD Xtreme Hot Start DNA Polymerase (Sigma-Aldrich) using the primers listed in the materials table. PCR products were run on 1.8% agarose gel to detect *eIF4G2* floxed (798 bp), wild type (730 bp) or excised (191 bp) bands.

### Histology and immunohistochemistry

Mouse organs were fixed in 4% paraformaldehyde (PFA) in phosphate-buffered saline (PBS) at 4°C overnight with gentle agitation. Specimens were washed 3 times in PBS, and incubated with 25%, 50% and 70% ethanol in PBS for 30 minutes each and kept in 70% ethanol at 4°C until further processed. Tissue dehydration, embedding, sectioning (4 μm), H&E staining, PAS staining, cleaved caspase 3 and PCNA immunohistochemistry, and imaging for these slides were performed at Histowiz Inc (NY, USA).

For immunofluorescence, intestinal tissue sections were de-waxed by immersion in xylene (5 min, 2 times) and hydrated by serial immersion in 100% EtOH (5 min, 2 times), 95% EtOH (5 min), 70% EtOH (3 min) and distilled water (3 min). Antigen retrieval was performed by heating sections in Target Retrieval Solution, Citrate pH 6.1 (DAKO) at 121°C for 20 min. After cooling down and a brief rinse with distilled water, sections were incubated with blocking buffer (PBS containing 5% donkey serum, 1% BSA, and 0.1% Tween20) for 1 hour at room temperature. Primary antibodies diluted in blocking buffer were then added and incubated overnight at 4°C. After washing 3 times in PBS containing 0.1% Tween20, sections were incubated with secondary antibodies diluted in blocking buffer for 1.5 hours at room temperature, protected from light. Nuclei were stained with DAPI. Sections were washed 3 times in PBS containing 0.1% Tween20 and mounted with Fluormount G mounting medium (Invitrogen) and a cover slip placed over the tissue section. Antibodies used were: rabbit monoclonal anti-lysozyme (1:200, ab108508, Abcam), rabbit multiclonal anti-chromogranin A (1:500, ab283265, Abcam), rabbit monoclonal anti-DCAMKL1 (1:50, ab109029, Abcam), rabbit anti-active YAP (1:200, ab205270, Abcam), Alexa 488-conjugated donkey anti-rabbit IgG (1:500, A-21206, Thermo Fisher Scientific), Alexa 647-conjugated donkey anti-rabbit IgG (1:500, A-31573, Thermo Fisher Scientific). Stained sections were imaged using the Keyence BZ-X810 microscope.

### Pathological evaluation of the small intestine

To assess the severity of lesions in the small intestine, a personalized scoring system was adapted from a previously published paper^29^. Slides were examined and scored by a board-certified veterinary pathologist blinded to the experimental groups during the initial evaluation. Overall, each mouse received a score according to a five-tier severity scale (0, absent; 1, minimal / <10% rare or infrequent finding; 2, mild / 10-25% noticeable but not prominent finding; 3, moderate / 25-50% prominent but not dominant finding; 4, marked / >50% dominating and/or overwhelming finding) for each of the following parameters: goblet cell depletion, Paneth cell depletion, crypt proliferation, crypt single cell necrosis, epithelial erosion, epithelial ulceration, crypt loss, villous blunting, inflammatory cell infiltration and inflammatory cell extension.

### RNAscope and BaseScope

RNA in situ hybridization was performed on paraffin-embedded tissue specimens using either RNAscope 2.5 High Definition-RED Assay or BaseScope Reagenet Kit v2 RED (Advanced Cell Diagnostics) following the manufacturer’s instructions. Briefly, after de-waxing and hydration, specimens were incubated with hydrogen peroxide reagent at RT for 10 min, target retrieval at 100°C for 15 min, followed by either protease plus (for RNAscope) or protease III (for BaseScope) treatment at 40°C for 30 min. RNAscope or BaseScope probes (see Materials Table) were hybridized at 40°C for 2 hours. Nuclei and tissue sections were counterstained with hematoxylin. At least 3 biological replicates were tested. Stained sections were imaged using a Keyence BZ-X810 microscope.

To quantify *Bmi1* RNA along the crypt-villus axis, we analyzed ten well-oriented crypts per mouse (three mice per genotype; 30 crypts per genotype). Crypt cell positions were annotated with the absolute coordinate method: the nucleus in contact with the basement membrane was designated position 0, and positions were incremented by one for each successive nucleus towards the villus tip, irrespective of cell type. Every epithelial cell in each crypt was scored for *Bmi1* signal intensity according to the RNAscope 2.5 HD-Red semi-quantitative guideline provided by Advanced Cell Diagnostics (0 = 0 dots /cell; 1 = 1–3 dots /cell; 2 = 4–9 dots /cell; 3 = 10–15 dots /cell; 4 ≥ 15 dots or >10 % clusters). Statistical analysis was performed in GraphPad Prism 10. Cell position and genotype were modelled with a mixed-effects two-way ANOVA (REML estimation) and Geisser–Greenhouse correction for possible sphericity violations. Sidák-adjusted pair-wise comparisons were applied to assess genotype differences at each cell position.

### Production of R spondin-conditioned media

R-spondin 1 expressing HEK293T (Sigma-Aldrich) cells were maintained in DMEM, 20% fetal calf serum, 1% penicillin/streptomycin, and 125 µg/mL Zeocin (Thermo Fisher Scientific) at 37°C, 5% CO_2_. Cells were cultured in organoid basal medium (Advanced DMEM/F12 (Gibco) supplemented with 10 mmol/l HEPES (Gibco), 1x Glutamax (Gibco), 1x B27 (Gibco), and 1 mM N-acetylcysteine (Sigma)) for 5 days and the supernatant was collected as R-spondin 1-conditioned media.

### Organoid establishment and culture

Mouse organoids were established and maintained from isolated crypts of the proximal small intestine as previously described^42^. Briefly, immediately after euthanizing with CO_2_ inhalation and cervical dislocation, the proximal half of the small intestine was harvested and flushed with PBS before being cut open longitudinally. The villi were removed by scraping with a microscope slide. Pieces of approximately 5 mm length were rotated in 2.5 mM EDTA in PBS for 20 min at 4°C. After removal of EDTA, tissue fragments were vigorously shaken in 10% FBS/PBS to release crypts. Supernatant fractions enriched in crypts were passed through a 40-μm cell strainer and centrifuged at 300g for 5 min. The cell pellet was washed with DMEM supplemented with antibiotics and counted under a microscope. After centrifugation at 300g for 5 min, crypts were resuspended in Matrigel (growth factor reduced, Corning) and plated in 12-well plates (200 crypts per 20 μl Matrigel dome, 4 domes per well). Following polymerization of Matrigel, 1 ml of ENR culture medium (Organoid basal medium supplemented with EGF (50 ng/ml, Gibco), Noggin (100 ng/ml, PeproTech) and 10% R-spondin conditioned media). Organoids were passaged every 4-5 days by mechanical disruption at 1:4 – 1:6 ratio, as previously described^68^. To activate CreER, organoids were treated with 0.5 μM 4-OHT (Sigma) for 24 hours. For differentiation assays, organoids were first treated with ENR medium supplemented with 5 μM CHIR99021 (Sigma) and 1 mM valproic acid (Sigma) for 6 days to induce ISC enrichment. On the 6^th^ day, DMSO or 0.5 uM 4-OHT was added to the medium. Organoids were then passaged at 1:6 ratio and differentiated by culturing in ENR medium supplemented with the following; 5 μM CHIR99021 and 10 uM DAPT (Tocris) for Paneth cell differentiation, 2 uM IWP-2 (Sigma) and 10 uM DAPT for goblet cell differentiation, and 2 uM IWP-2 and 1 mM valproic acid for enterocyte differentiation. 10% Afamin/Wnt3a conditioned medium (MBL International) was added to the ENR medium where indicated (WENR condition). For CREBBP/EP300 inhibitor experiments, A-485 (Tocris, 3-10 μM) or DMSO was added to ENR medium for 4 days.

### Organoid whole-mount staining

Organoid whole-mount staining was performed as previously described^68^. Organoids were incubated with anti-active YAP antibody (abcam) overnight at 4°C, and then stained with Alexa 488-conjugated donkey anti-rabbit IgG (1:500, A-21206, Thermo Fisher Scientific) and DAPI. Organoids were imaged using a Zeiss LSM880 confocal microscope.

### RNA isolation and RT-qPCR analysis

Intestinal crypt and villi samples were collected as described above. After washing with ice-cold PBS, 350ul of RLT plus reagent (QIAGEN) was added to 10mg of tissue and were lysed using TissueLyser II (QIAGEN). RNA from tissue was extracted using RNeasy mini kit (QIAGEN) with on-column DNA digestion with RNase-Free DNase (QIAGEN). Organoids were recovered from Matrigel domes by incubating in Cell Recovery Solution (Corning) at 4°C on a shaker for 1 hour. After washing with ice-cold PBS, organoids were lysed and total RNA was purified using a RNeasy micro kit (QIAGEN) with on-column DNA digestion with RNase-Free DNase (QIAGEN). Purified RNA (0.1-1ug) was used for single stranded complementary DNA (cDNA) synthesis using PrimeScript RT Master Mix (Takara Bio). Quantitative RT-PCR was performed using TaqMan Fast Advanced Master Mix (Applied Biosystems) on either StepOne instrument (Applied Biosystems) or QuantStudio 5 (Applied Biosystems). All target probes (FAM-MGB) were multiplexed with mouse *Gapdh* probe (VIC-MGB_PL) within the well. The levels of mRNA were normalized to mouse *Gapdh* expression, and then relative expression was calculated as the fold-change from the control. Primer information is provided in the Materials Table.

### RNA sequencing

To compare gene expression between *Eif4g2*^+/-^ and *Eif4g2*^-/-^ intestinal crypts and villi, *Eif4g2*^flox/+^; *Rosa26*-CreER and *Eif4g2*^flox/flox^; *Rosa26*-CreER mice were treated with Tamoxifen as above and dissected at day 9 after the first administration. To compare gene expression in SIOs (ENR and WENR condition), *Eif4g2*^+/+^ and *Eif4g2*^flox/flox^ SIOs were first cultured in ENR or WENR medium for 6 days. From the day of passaging, SIOs were treated with ENR or WENR medium containing DMSO or 0.5μM 4-OHT for 24 hours. Samples were collected after being cultured for another 5 days in ENR or WENR medium. For RNA sequencing of A-485 treated SIOs, cells were maintained in ENR medium. From the day of passaging, SIOs were treated with either DMSO, 3, 5, or 10μM of A-485 for 4 days and then collected. RNA was extracted as above and subsequently sent to Novogene (Sacramento, CA) for library preparation, sequencing (Illumina, PE150, 30M paired reads) and bioinformatics analyses, including adapter trimming and quality control using their in-house scripts. Reads were aligned to the mouse reference genome (GRCm38/mm10, Genome Reference Consortium, downloaded from https://www.ncbi.nlm.nih.gov/assembly/GCF_000001635.20/) using Hisat2^69^ v2.0.5. Gene expression quantification was performed using featureCounts^70^ v1.5.0-p3 with options -s 2 to account for strand-specificity. Differential expression analysis was performed using DESeq2^71^ v1.20.0. P-values were adjusted using the Benjamini and Hochberg’s approach for controlling the false discovery rate. Genes with an adjusted P-value ≤ 0.05 were assigned as differentially expressed. Gene ontology analyses was performed using EnrichR^72–74^. Gene set enrichment analyses was performed using clusterProfiler^75^. The fetal spheroid signature gene set was based on genes upregulated >2 fold in fetal tissue derived spheroids versus adult tissue derived organoids^36^. The YAP-activated gene signatures were curated from the literature describing YAP activated genes in intestinal cells^37,38^. Genes upregulated in *Eif4g2*^-/-^ SIOs were defined as genes with >2-fold change in the SIO RNA-seq (ENR condition).

### Library preparation for Thor-Ribo-seq and paired RNA-seq

For Ribo-seq and paired RNA-seq experiments, *Eif4g2*^+/+^, *Eif4g2*^fl/+^ and *Eif4g2*^fl/fl^ SIOs were maintained in ENR medium. On the day of passage, organoids were treated with 0.5 μM 4-OHT for 24 hours and cultured in ENR medium for another 3 days. Matrigel domes containing organoids were scraped off using spatulas and snap frozen in liquid nitrogen. The library of Thor-Ribo-Seq was prepared as previously described with modifications^76,77^. Twelve frozen domes containing organoids were combined with frozen droplets of 300 μl lysis buffer (20 mM Tris–HCl pH 7.5, 150 mM NaCl, 5 mM MgCl_2_, 1 mM dithiothreitol, 100 μg/ml cycloheximide, 100 μg/ml chloramphenicol, and 1% Triton X-100) and pulverized using Multi-beads Shocker (Yasui Kikai) at 2,800 rpm for 15 sec. Lysates were incubated with 15 U of Turbo DNase (Thermo Fisher Scientific) for 10 min on ice and clarified by centrifugation at 20,000 g for 10 min at 4°C. Lysates containing ∼2 μg of RNA were treated with 20 U of RNase I (LGC Biosearch Technologies) for 45 min at 25°C. Ribosomes were precipitated by 1 M sucrose cushion followed by ultracentrifugation for 1 h at 100,000 rpm (408,800 × g) at 4 °C using Optima MAX-TL and a TLA-110 rotor (Beckman Coulter). RNAs ranging from 17 to 34 nt were excised from 15% TBE–urea gels, purified, and ligated with DNA linkers containing the T7 promoter sequence. After the rRNA depletion using the riboPOOL rRNA Depletion Kit (Mouse-Rat Ribo-Seq) (siTOOLs Biotech), an oligonucleotide to the T7 promoter region was hybridized and in vitro transcription was performed using a T7-Scribe Standard RNA IVT Kit (CELLSCRIPT). Amplified RNAs were subsequently ligated with the second linker, reverse-transcribed, and PCR-amplified.

For paired RNA-seq, total RNA was purified using TRIzol LS reagent (Thermo Fisher Scientific) and Direct-zol RNA Microprep Kit (Zymo research). Following the removal of rRNA by riboPOOL Mouse-Rat rRNA depletion kit (siTOOLs Biotech), the sequencing library was prepared with SEQuoia Express Stranded RNA Library Prep Kit (Bio-Rad). The libraries were sequenced on a NovaSeq X Plus platform (Illumina) with a pair-end 150 bp read.

Data processing was performed as previously described^76^ with modifications. For Thor-Ribo-Seq, after the base correction with overlap analysis, the quality filtering and the adapter removal were performed on read 1 using fastp^78^. For RNA-seq, both paired reads were used. Reads were then demultiplexed by Cutadapt 3.7^79^ and mapped to noncoding RNAs using STAR v2.7.0a^80^ to exclude them from the analysis. The remaining reads were aligned to the mouse genome mm39 and assigned to the GENCODE Mouse release VM36 reference obtained from the UCSC Genome Browser (https://genome.ucsc.edu/index.html) using STAR. UMI suppression was performed using UMItools v1.1.2^81^. Read counting was performed using the custom scripts (https://github.com/ingolia-lab/RiboSeq). The offsets of the A site from the 5′ end of the ribosome footprint were determined to be 15 for fragments of 19–34 nt. For RNA-Seq, an offset of 15 was used for all mRNA fragments. The canonical transcripts were defined based on Ensembl Canonical.

The foldchange calculations were performed with the DESeq2 package^71^. Translation efficiency (TE), which are Ribo-Seq counts normalized by RNA-seq counts, was also measured by DESeq2. The significance was calculated by a likelihood ratio test in a generalized linear model. The reads corresponding to the first and last five codons in the coding sequences were excluded from the calculation. mRNAs were defined as uORF-containing if they included at least one uORF annotated in Ribo-uORFs with a strong Kozak sequence. Gene ontology analyses were performed using EnrichR^72–74^. All custom scripts used in this study are available upon request.

### Protein extraction

Mouse intestinal crypts and organoids were isolated as above and washed in ice-cold PBS. 500 ul RIPA buffer (Thermo Scientific) supplemented with 1% protease inhibitors and phosphatase inhibitors (Sigma-Aldrich) was added to approximately 10 mg of isolated intestinal crypts and samples were lysed by applying to TissueLyser II (QIAGEN). Organoids were lysed with ice-cold lysis buffer (either MPER (Thermo Scientific) or Laemmli buffer (Bio-Rad), depending on the downstream targets) with 1% phosphatase and protease inhibitors. After removing undissolved debris by centrifugation, protein concentration was determined, and samples were snap frozen and stored at -80°C until analysis.

### Protein expression analysis by antibody-based protein quantification

Antibody-based protein quantification was performed using a WES/Jess automated capillary electrophoresis system (proteinsimple) with 12-230 kDa or 66-440 kDa Separation Module (proteinsimple) as instructed. The protein concentration was adjusted to 0.4 or 0.8 mg/mL depending on the target protein. Target protein expression was normalized either by multiplexing with the GAPDH or Histone H3 antibody, or by using the RePlex module and total protein assay kit (both from proteinsimple). Antibodies used were: Mouse monoclonal anti-eIF4G2 (1:1000, 610742, BD Biosciences), rabbit monoclonal anti-OLFM4 (1:100, 14369S, Cell Signaling Technologies), rabbit monoclonal anti-lysozyme (1:50, ab108508, Abcam), rabbit anti-active YAP (1:200, ab205270, abcam), rabbit monoclonal anti-acetyl-histone H3 (Lys18) (1:10, 13998S, Cell Signaling Technologies), rabbit monoclonal anti-acetyl-histone H3 (Lys27) (1:10, 8173S, Cell Signaling Technologies), rabbit monoclonal anti-monomethyl-histone H3 (Lys4) (1:10, 5326S, Cell Signaling Technologies), rabbit monoclonal anti-histone H3 (1:200, 4499T, Cell Signaling Technologies), rabbit polyclonal anti-GAPDH (1:100, ab9485, Abcam). Data was analyzed and visualized using Compass for Simple Western software (Protein Simple). The WES protein images were used throughout the manuscript.

### Proteomics

Organoids were lysed in PTS buffer (12 mM SDC, 12 mM SLS, and 100 mM Tris-HCl (pH 9.0)) containing a protease and phosphatase inhibitor mixture (Sigma-Aldrich). The lysates were used as protein samples and processed using a modified protein aggregation capture method^82^ as previously described^83^. A total of 200 ng of desalted peptides were loaded and separated on an Aurora column (250 mm length, 75 mm i.d., IonOpticks) using a nanoElute (Bruker) for subsequent analysis by timsTOF Pro2 system (Bruker). The mobile phases were composed of 0.1% formic acid (solution A) and 0.1% formic acid in acetonitrile (solution B). A flow rate of 400 nL/min of 2-17% solution B for 60 min, 17-25% solution B for 30 min, 25-37% solution B for 10 min, 37-80% solution B for 10 min, and 80% solution B for 10 min was used (120 min in total). The applied spray voltage was 1400 V, and the interface heater temperature was 180°C. To obtain MS and MS/MS spectra, the Parallel Accumulation Serial Fragmentation (PASEF) acquisition method with data-independent acquisition (DIA) mode was used (diaPASEF)^84^. For diaPASEF settings, 1.7 s per one cycle with precursor ion scan and 16 times diaPASEF scans were conducted with the MS/MS isolation width of 25 m/z, precursor ion ranges of 400-1200 m/z, ion mobility ranges of 0.57-1.47 V・s・cm(-2). The obtained DIA data were searched by DIA-NN (v1.8.2 beta27)^85^ against selected mouse entries of UniProt/Swiss-Prot release 2022_03 with the carbamidomethylation of cysteine as the fixed modification and protein N-terminal acetylation and methionine oxidation as the variable modification. For the other DIA-NN parameters, Trypsin/P protease, one missed cleavage, peptide length range of 7-30, precursor m/z range of 300-1800, precursor charge range of 1-4, fragment ion m/z range of 200-1800, and 1% precursor FDR were used. The values ‘‘PG.MaxLFQ’’ from the results were used as representative protein area values for the comparison of protein expression levels.

### Quantification and statistical analysis

Data in the figure panels are presented as mean ± standard deviation. Sample number (n) indicates the number of biological replicates in each experiment. For RT-qPCR and Wes data, comparisons between two groups were evaluated using two-tailed, unpaired Student’s t-tests. To compare intestinal pathological scores within different groups, Kruskal-Wallis test was performed. Comparison of body weight change in mice was done using two-way ANOVA with repeated measures to assess the effects of genotype (+/+, +/-, -/-), time (days or weeks post-tamoxifen injection), and their interaction on body weight changes. Tukey’s post-hoc test was applied for multiple comparisons between genotypes at each time point. All analyses were performed using GraphPad Prism 10 software (San Diego, CA, USA). Significant differences are noted at p < 0.05, and range of statistical significance is shown by an asterisk within the figure panels: ns = not significant, ^∗^ = p < 0.05, ^∗∗^ = p < 0.01, ^∗∗∗^ = p < 0.001, ^∗∗∗∗^ = p < 0.0001.

## Data availability

Thor-Ribo-Seq (GSE294619) and RNA-seq data (GSE294512) obtained in this study were deposited in the National Center for Biotechnology Information (NCBI) database. The mass spectrometry data were deposited in the ProteomeXchange Consortium via jPOSTrepo^86^ (https://repository.jpostdb.org/) with the dataset identifier JPST003517 (PXD059198).

## Code availability

Custom scripts for the mapping and quantification for ribosome profiling are available at https://github.com/ingolia-lab/RiboSeq.

