## Supplemental figures 1-3, supplmental table 1-2, supplmental methods for "eIF4G2-Mediated Translation Initiation of Histone Modifiers Is Essential for Intestinal Stem Cell Maintenance and Differentiation"

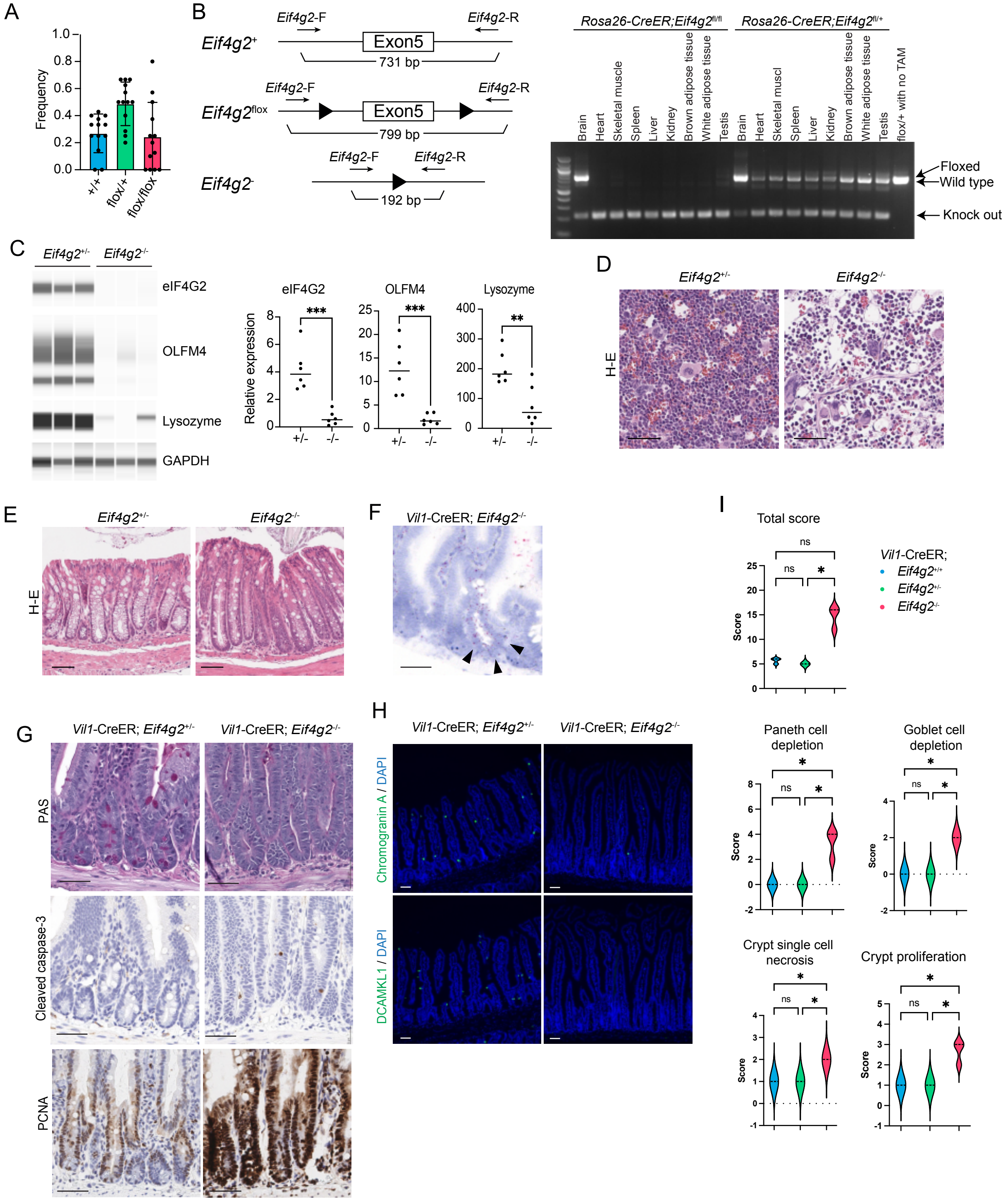


Supplemental Figure 1.

Supplemental Figure 1. (A) Frequency of genotype in offsprings from intercrosses between *Eif4g2*$^{flox/+}$ mice. (B) PCR result of the *Eif4g2* locus of various organs from the *Rosa26*-CreER; *Eif4g2*$^{flox/flox}$ and *Eif4g2*$^{flox/+}$ mice 14 days after tamoxifen treatment. (C) Protein expression analyses of intestinal crypts obtained from *Rosa26*-CreER*; Eif4g2*$^{+/-}$ and *Eif4g2*$^{-/-}$ mice at endpoint. (Left) Wes run image, (Right) quantification of proteins normalized to GAPDH expression. n=6 each. Bars indicate the mean of the replicates. (D) H-E stain of *Rosa26*-CreER; *Eif4g2*$^{+/-}$ and *Eif4g2*$^{-/-}$ mouse bone marrow 14 days after tamoxifen treatment. (E) H-E stain of *Rosa26*-CreER; *Eif4g2*$^{+/-}$ and *Eif4g2*$^{-/-}$ mouse colon 14 days after tamoxifen treatment. (F) *Eif4g2* mRNA BaseScope in *Vil1*-CreER; *Eif4g2*$^{flox/flox}$ mice one month post tamoxifen treatment indicating a crypt that escaped the gene KO (arrowhead). (G) Histological findings in *Vil1*-CreER; *Eif4g2*$^{+/-}$ and *Eif4g2*$^{-/-}$ mice 1 month after tamoxifen treatment. From top; PAS staining, cleaved caspase-3 IHC, PCNA IHC. (H) Fluorescent IHC of *Vil1*-CreER; *Eif4g2*$^{+/-}$ and *Eif4g2*$^{-/-}$ mice 1 month after tamoxifen treatment. Top; enteroendocrine cell marker chromogranin A, Bottom; tuft cell marker DCAMKL1. (I) Violin plots showing modified intestinal lesion score of *Vil1*-CreER; *Eif4g2*$^{+/+}$, *Eif4g2*$^{+/-}$ and *Eif4g2*$^{-/-}$ mice 1 month after tamoxifen treatment (n=3 each. See also Table S2). * p <0.05, ** p < 0.005, *** p < 0.001. Scale bars indicate 50 um.

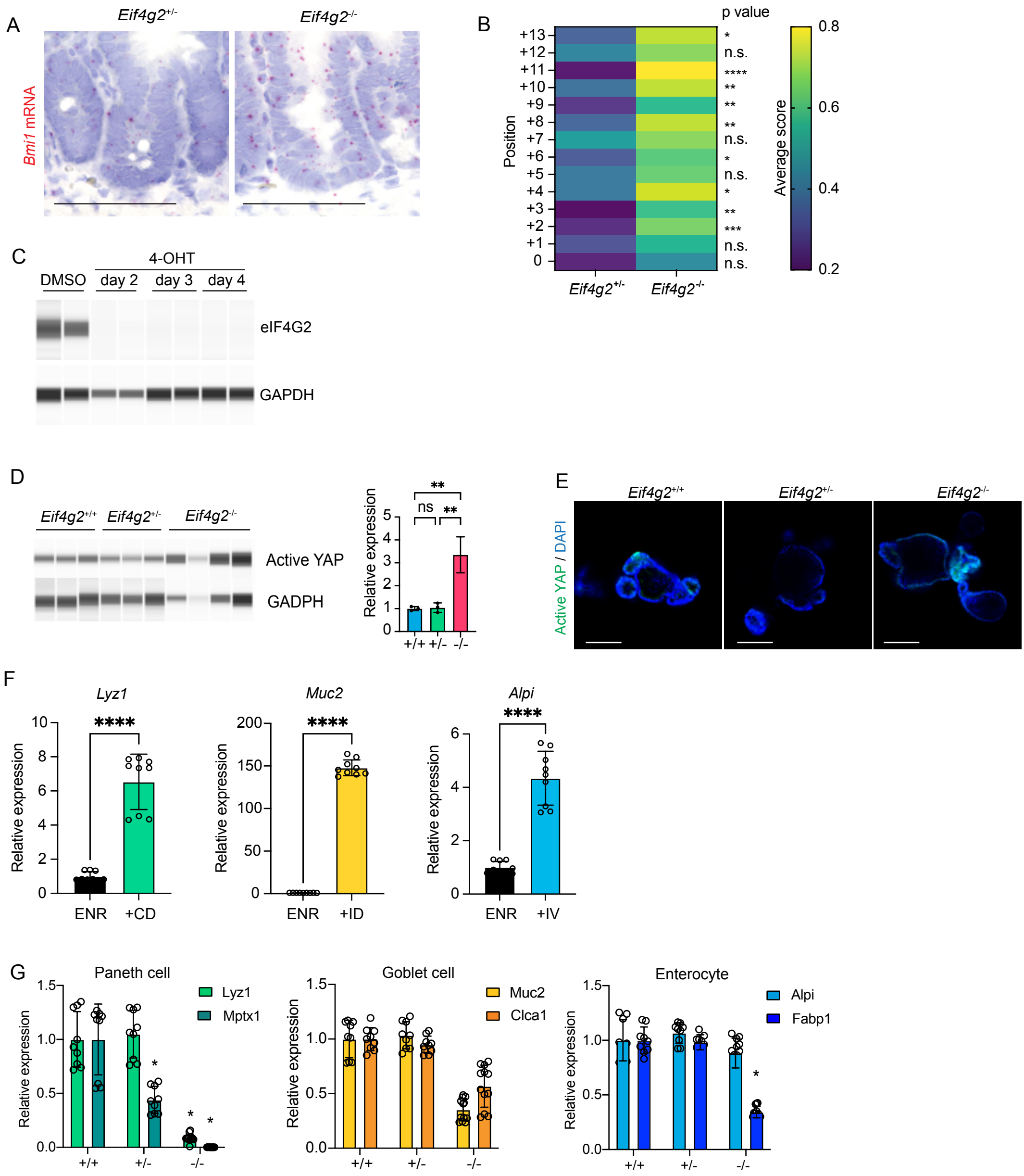


Supplemental Figure 2.

Supplemental Figure 2. (A) *Bmi1* RNAscope results in *Rosa26-CreER*; *Eif4g2*$^{+/-}$ and *Eif4g2*$^{-/-}$ intestinal crypts. (B) Heatmap showing the semi-quantitative scores of *Bmi1* expression in *Eif4g2*$^{+/-}$ and *Eif4g2*$^{-/-}$ intestinal crypts. Ten crypts from three mice in each genotype were examined. (C) Wes run image of eIF4G2 and GAPDH protein expression in *Eif4g2*$^{flox/flox}$; *Rosa26*-CreER SIOs treated with DMSO and 2, 3, and 4 days after 4-OHT treatment. (D) Protein expression analyses of Active YAP in *Eif4g2*$^{+/+}$, *Eif4g2*$^{+/-}$ and *Eif4g2*$^{-/-}$ SIOs 20 days after 4-OHT treatment. (Left) Wes run image, (Right) quantification of proteins normalized to GAPDH expression. n=4 each. (E) Whole-mount active YAP staining of *Eif4g2*$^{+/+}$, *Eif4g2*$^{+/-}$ and *Eif4g2*$^{-/-}$ SIOs 10 days after 4-OHT treatment. (F) RT-qPCR analysis in wild type SIOs treated with (left) ENR and ENR supplemented with CHIR99021 and DAPT (+CD), (middle) ENR and ENR supplemented with IWP-2 and DAPT (+ID), (right) ENR and ENR supplemented with IWP-2 and Valproic acid (+IV). (G) RT-qPCR analysis of *Eif4g2*$^{+/+}$, *Eif4g2*$^{+/-}$ and *Eif4g2*$^{-/-}$ SIOs treated with (left) CHIR99021 and DAPT for Paneth cells, (middle) IWP-2 and DAPT for goblet cells, (right) IWP-2 and valproic acid for enterocytes. n=3, 3 technical replicates each. Data are represented as mean +/- SEM. * $p <0.05$, ** $p < 0.005$, *** $p < 0.001$, **** $p < 0.0001$. Scale bars indicate 50 um.

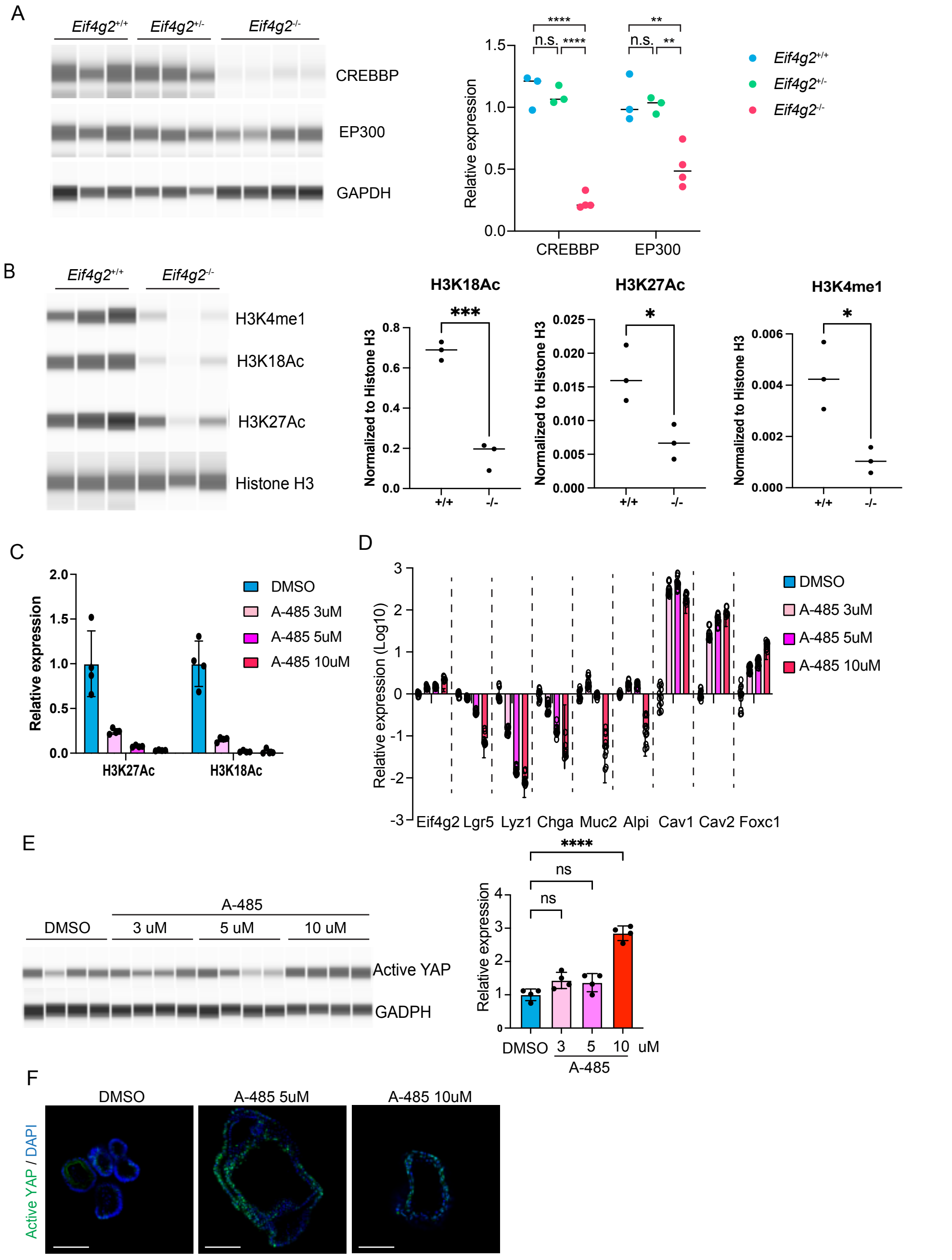


Supplemental Figure 3

Supplemental Figure 3. (A) Protein quantification of CREBBP, EP300 and GAPDH in *Eif4g2*$^{+/+}$, *Eif4g2*$^{+/-}$ and *Eif4g2*$^{-/-}$ SIOs 6 days after 4-OHT treatment. (Left) Wes run image, (Right) quantification of proteins normalized to GAPDH expression. n=3 for +/+ and +/-, n=4 for -/-. Bars indicate the mean of the replicates. (B) Protein quantification of H3K4me1, H3K18ac, H3K27ac, and Histone H3 in *Eif4g2*$^{+/+}$ and *Eif4g2*$^{-/-}$ SIOs 18 days after 4-OHT treatment (n=3 each). (Left) Wes run image, (Right) quantification of proteins normalized to Histone H3 expression. Bars indicate the mean of the replicates. (C) Protein quantification of H3K18ac and H3K27ac in SIOs with DMSO or A-485 (3, 5, or 10uM, left) for 4 days. n=4 each. Protein expression is normalized to total protein amount. (D) RT-qPCR analysis of SIOs treated with DMSO or A-485 (3, 5, or 10uM) for 4 days. n=4 each. Data are represented as mean +/- SEM. (E) Protein quantification of active YAP normalized to GAPDH in SIOs treated with DMSO or A-485 (3, 5, or 10uM) for 4 days. (Left) Wes run image, (Right) quantification of proteins normalized to GAPDH expression. n=4 each. (F) Whole-mount active YAP staining of SIOs treated with DMSO or A-485 (5, 10uM) for 4 days. Data are represented as mean +/- SEM. * $p < 0.05$, ** $p < 0.005$, *** $p < 0.001$, **** $p < 0.0001$. Scale bars indicate 100um.

Table S1. Modified intestinal lesion score of *Eif4g2*$^{+/-}$ and *Eif4g2*$^{-/-}$ (*Rosa26*-CreER) small intestines at end point

| | | Epithelial changes | | | | Mucosal architecture | | | | Inflammation | | |
|---|---|---|---|---|---|---|---|---|---|---|---|---|
| Animal ID | Experimental group | Goblet cell depletion | Paneth cell depletion | Crypt proliferation | Crypt single cell necrosis/apoptosis | Epithelial Erosion | Epithelial Ulceration | Crypt loss | Villous blunting | Inflammatory grade (severity) | Inflammatory extend (layers) | TOTAL SCORE (SUM) |
| 4384 | Hetero-KO | 0 | 0 | 1 | 1 | 0 | 0 | 0 | 0 | 1 | 1 | 4 |
| 4385 | Homo-KO | 2 | 2 | 2 | 2 | 0 | 0 | 0 | 0 | 1 | 1 | 10 |
| 4386 | Homo-KO | 0 | 1 | 1 | 1 | 0 | 0 | 0 | 0 | 1 | 1 | 5 |
| 4389 | Hetero-KO | 0 | 0 | 1 | 1 | 0 | 0 | 0 | 0 | 1 | 1 | 4 |
| 4291 | Hetero-KO | 0 | 0 | 0 | 0 | 0 | 0 | 0 | 1 | 1 | 1 | 3 |
| 4292 | Hetero-KO | 0 | 0 | 0 | 0 | 0 | 0 | 0 | 1 | 1 | 1 | 3 |
| 4315 | Homo-KO | 1 | 1 | 1 | 1 | 0 | 0 | 0 | 0 | 1 | 1 | 6 |
| 4316 | Homo-KO | 2 | 3 | 3 | 2 | 0 | 0 | 0 | 0 | 1 | 1 | 12 |
| 6040 | Hetero-KO | 0 | 0 | 0 | 0 | 0 | 0 | 0 | 1 | 1 | 1 | 3 |
| 6041 | Homo-KO | 0 | 0 | 0 | 1 | 0 | 0 | 0 | 1 | 1 | 1 | 4 |
| 6042 | Homo-KO | 3 | 4 | 3 | 2 | 0 | 0 | 0 | 1 | 3 | 2 | 18 |
| 6043 | Hetero-KO | 0 | 0 | 0 | 0 | 0 | 0 | 0 | 0 | 1 | 1 | 2 |
| 6078 | Hetero-KO | 0 | 0 | 1 | 0 | 0 | 0 | 0 | 1 | 1 | 1 | 4 |
| 6079 | Homo-KO | 2 | 4 | 3 | 2 | 0 | 0 | 0 | 1 | 2 | 2 | 16 |
| 7840 | Hetero-KO | 0 | 0 | 0 | 0 | 0 | 0 | 0 | 1 | 1 | 1 | 3 |
| 7842 | Homo-KO | 1 | 1 | 1 | 1 | 0 | 0 | 0 | 1 | 1 | 1 | 7 |
| 7843 | Hetero-KO | 0 | 0 | 0 | 0 | 0 | 0 | 0 | 1 | 1 | 1 | 3 |
| 7871 | Hetero-KO | 0 | 0 | 1 | 0 | 0 | 0 | 0 | 1 | 1 | 1 | 4 |
| 7872 | Hetero-KO | 0 | 0 | 1 | 0 | 0 | 0 | 0 | 1 | 1 | 1 | 4 |
| 7909 | Hetero-KO | 0 | 0 | 1 | 0 | 0 | 0 | 0 | 1 | 1 | 1 | 4 |
| 7910 | Hetero-KO | 0 | 0 | 1 | 0 | 0 | 0 | 0 | 1 | 1 | 1 | 4 |
| 7911 | Homo-KO | 3 | 3 | 3 | 2 | 0 | 0 | 0 | 1 | 2 | 2 | 16 |
| 7921 | Homo-KO | 2 | 2 | 2 | 2 | 0 | 0 | 0 | 1 | 1 | 1 | 11 |
| 7924 | Homo-KO | 3 | 2 | 3 | 2 | 0 | 0 | 0 | 1 | 2 | 2 | 15 |

Table S2. Modified intestinal lesion score of *Vil1*-CreER; *Eif4g2*$^{+/-}$ and *Eif4g2*$^{-/-}$ small intestines 1 month after tamoxifen treatment

| | | Epithelial changes | | | | Mucosal architecture | | | | Inflammation | | TOTAL SCORE (SUM) |
|---|---|---|---|---|---|---|---|---|---|---|---|---|
| Animal ID | SIMPLE GROUS | Goblet cell depletion | Paneth cell depletion | Crypt proliferation | Crypt single cell necrosis/apoptosis | Epithelial Erosion | Epithelial Ulceration | Crypt loss | Villous blunting | Inflammatory grade (severity) | Inflammatory extend (layers) | |
| 6574 | Wild type | 0 | 0 | 1 | 1 | 0 | 0 | 0 | 2 | 1 | 1 | 6 |
| 6689 | Wild type | 0 | 0 | 1 | 1 | 0 | 0 | 0 | 2 | 1 | 1 | 6 |
| 6693 | Wild type | 0 | 0 | 1 | 1 | 0 | 0 | 0 | 1 | 1 | 1 | 5 |
| 6690 | Hetero-KO | 0 | 0 | 1 | 1 | 0 | 0 | 0 | 1 | 1 | 1 | 5 |
| 6695 | Hetero-KO | 0 | 0 | 1 | 1 | 0 | 0 | 0 | 1 | 1 | 1 | 5 |
| 6698 | Hetero-KO | 0 | 0 | 1 | 1 | 0 | 0 | 0 | 1 | 1 | 1 | 5 |
| 6692 | Homo-KO | 2 | 4 | 3 | 2 | 0 | 0 | 0 | 2 | 2 | 1 | 16 |
| 6696 | Homo-KO | 2 | 4 | 3 | 2 | 0 | 0 | 0 | 2 | 2 | 1 | 16 |
| 6697 | Homo-KO | 2 | 2 | 2 | 2 | 0 | 0 | 0 | 1 | 2 | 1 | 12 |

**Materials**

| Reagent or Resource | Source | Identifier |
|---|---|---|
| **Mouse lines** | | |
| *Eif4g2*$^{flox}$ | This paper | N/A |
| B6.129-*Gt(ROSA)26Sor*$^{tm1(cre/ERT2)Tyj}$/J (ROSA26-CreERT2) | Jackson Laboratories | #008463 |
| B6.Cg-Tg(Vil1-cre/ERT2)23Syr/J (Villin-CreERT2) | Jackson Laboratories | #020282 |
| **Antibodies** | | |
| Mouse monoclonal anti-NAT1 | BD Biosciences | 610742, RRID:AB_398065 |
| Rabbit monoclonal anti-OLFM4 | Cell Signaling Technology | 14369S, RRID: AB_2798465 |
| Rabbit monoclonal anti-Lysozyme | Abcam | Ab108508,RRID:AB_10861277 |
| Rabbit multiclonal anti-Chromogranin A | Abcam | Ab283265 |
| Rabbit monoclonal anti-DCMKL1 | Abcam | Ab109029, RRID:AB_10864128 |
| Rabbit polyclonal anti-cleaved caspase-3 | Cell Signaling Technology | 9661,RRID:AB_2341188 |
| Rabbit monoclonal anti-PCNA | Cell Signaling Technology | 13110,RRID:AB_2636979 |
| Recombinant Rabbit anti-active YAP | abcam | Ab205270, RRID: AB_2813833 |
| Rabbit polyclonal anti-GAPDH | Abcam | Ab9485, RRID:AB_307275 |
| Rabbit monoclonal anti-Histone H3 | Cell Signaling Technology | 4499T |
| Rabbit monoclonal anti-acetyl-histone H3 (Lys18) | Cell Signaling Technology | 13998S |
| Rabbit monoclonal anti-monomethyl-histone H3 (Lys4) | Cell Signaling Technology | 5326S |
| Rabbit monoclonal anti-acetyl-histone H3 (Lys27) | Cell Signaling Technology | 8173S |
| Donkey anti-Rabbit IgG (H+L) Highly Cross-Adsorbed Secondary Antibody, Alexa Fluor 488 | Invitrogen | A-21206, RRID: AB_2535792 |
| Donkey anti-Rabbit IgG (H+L) Highly Cross- | Invitrogen | A-31573, RRID: AB_2536183 |

| | | |
|---|---|---|
| Adsorbed Secondary Antibody, Alexa Fluor 647 | | |
| **Chemicals, peptide, recombinant proteins** | | |
| Tamoxifen | Sigma-Aldrich | T5648 |
| (Z)-4-Hydroxytamoxifen | Sigma-Aldrich | H7904 |
| Fluoromount G mounting medium | Invitrogen | 00-4958-02 |
| DirectPCR Lysis Reagent (Tail) | Viagen Biotech | 102-T |
| Proteinase K solution | Viagen Biotech | 501-PK |
| KOD Xtreme Hot Start DNA Polymerase | Sigma-Aldrich | 71975 |
| Advanced DMEM/F12 | Gibco | 12634028 |
| B-27 Supplement | Gibco | 17504044 |
| HEPES | Gibco | 15630106 |
| GlutaMAX | Gibco | 35050-061 |
| N-acetyl-L-cysteine | Sigma-Aldrich | A9165-5G |
| Penicillin/Streptomycin solution | Corning | 30-002-CI |
| Zeocin | Gibco | R25001 |
| Matrigel Growth Factor Reduced Basement Membrane Matrix, Phenol Red-free, LDEV-free | Corning | 356231 |
| Afamin/Wnt3a-conditioned medium | MBL International | J2-001 |
| Mouse recombinant EGF | Invitrogen | PMG8043 |
| Recombinant human Noggin | Peprotech | 120-10C-250UG |
| CHIR99021 | Sigma-Aldrich | SML1046-5MG |
| Valproic acid | Sigma-Aldrich | P4543-25G |
| DAPT | Tocris | 2634/10 |
| IWP-2 | Sigma-Aldrich | I0536-5MG |
| A-485 | Tocris | 6387 |
| Cell Recovery Solution | Corning | CB-40253 |
| 1M Tris-HCl pH 7.5 | Fujifilm Wako Pure Chemical | 318-90225 |
| 1M Magnesium Chloride Solution | Nacalai tesque | 20942-34 |
| USB Dithiothreitol (DTT), 0.1M solution | Thermo Fisher Scientific | 707265ML |
| Cycloheximide | Merck | C4859-1ML |
| Triton X-100 | Nacalai tesque | 12967-32 |
| Turbo DNase | Thermo Fisher Scientific | AM2238 |

| | | |
|---|---|---|
| RNase I | LGC Biosearch Technologies | N6901K |
| TRIzol LS reagent | Thermo Fisher Scientific | 15596018 |
| MPER | Thermo Scientific | 78501 |
| Pierce RIPA buffer | Thermo Scientific | 89901 |
| 1M Tris-HCl pH9.0 | TEKNOVA | T1090 |
| Sodium deoxycholate | Sigma-Aldrich | 30970 |
| N-Lauroylsarcosine sodium salt | Sigma-Aldrich | 61743 |
| 2X Laemmli sample buffer | Bio-Rad | 1610737 |
| Protease inhibitor | Sigma-Aldrich | P8340-1ML |
| Phosphatase inhibitor cocktail 2 | Sigma-Aldrich | 5726-1ML |
| Phosphatase inhibitor cocktail 3 | Sigma-Aldrich | P0044-1ML |
| Target Retrieval Solution, Citrate pH 6.1 | Dako | S169984-2 |
| Alt-R S.p. Cas9 Nuclease V3 | Integrated DNA technologies | 1081058 |
| **Critical commercial assays** | | |
| Anti-Rabbit Detection Module for Jess, Wes, Peggy Sue or Sally Sue | proteinsimple | DM-001 |
| Anti-Mouse Detection Module for Jess, Wes, Peggy Sue or Sally Sue | proteinsimple | DM-002 |
| 20X Anti-Rabbit HRP Conjugate | proteinsimple | 043-426 |
| 12-230kDa Jess or Wes Separation Module, 8 x 25 capillary cartridges | proteinsimple | SM-W004 |
| 66-440kDa Jess or Wes Separation Module, 8 x 25 capillary cartridges | proteinsimple | SM-W008 |
| RePlex Module | proteinsimple | RP-001 |
| Total Protein Detection Module for Chemiluminescence based total protein assays | proteinsimple | DM-TP01 |
| RNeasy Mini Kit | QIAGEN | 74104 |
| RNeasy Plus Micro Kit | QIAGEN | 74034 |
| RNase-free DNase | QIAGEN | 79256 |
| PrimeScript RT Master Mix | Takara Bio | RR036A |
| Taqman Fast Advanced Master Mix | Applied Biosystems | 4444557 |

| | | |
|---|---|---|
| riboPOOL rRNA Depletion Kit (Mouse-Rat Ribo-Seq) | siTOOLs Biotech | dp-P012-052 |
| T7-Scribe Standard RNA IVT Kit | CELLSCRIPT | C-AS2607 |
| SEQuoia Express Stranded RNA Library Prep Kit | Bio-Rad | 12017297 |
| Direct-zol RNA Microprep Kit | Zymo Research | R2062 |
| Aurora column (250mm length, 75mm i.d.) | IonOptics | AUR3-25075C18-CSI |
| nanoElute | Bruker | NA |
| BaseScope Reagent Kit v2-Red Assay | Advanced Cell Diagnostics | 323900 |
| BaseScope Probe - BA-Mm-Eif4g2-2zz-st-C1 | Advanced Cell Diagnostics | 1090831-C1 |
| RNAscope 2.5 High Definition (HD)-RED Assay | Advanced Cell Diagnostics | 322350 |
| RNAscope Probe Mm-Bmi1-O1 | Advanced Cell Diagnostics | 466021 |
| RNAscope Probe Mm-Lgr5 | Advanced Cell Diagnostics | 312171 |
| **Oligonucleotides** | | |
| Mouse *Eif4g2* crRNA #1 | Integrated DNA technologies | TCAACTGATACTCTCCGGTG |
| Mouse *Eif4g2* gRNA #2 | Integrated DNA technologies | AACCAAATGGTTCACCCCCT |
| Alt-R CRISPR-Cas9 tracrRNA | Integrated DNA technologies | 1072532 |
| Mouse *Eif4g2* F primer | Integrated DNA technologies | GGTCTACAGAGTGAGTTCCAGGACAGC |
| Mouse *Eif4g2* R primer | Integrated DNA technologies | GCTCAAGCCAGACAAAAGCCCTACTCC |
| Eif4g2 (Mm00469038_m1) TaqMan Assay | Thermo Fisher Scientific | 4331182 |
| Lgr5 (Mm00438890_m1) TaqMan Assay | Thermo Fisher Scientific | 4331182 |
| Lyz1 (Mm00657323_m1) TaqMan Assay | Thermo Fisher Scientific | 4331182 |
| Mptx2 (Mm01621060_s1) TaqMan Assay | Thermo Fisher Scientific | 4331182 |
| Muc2 (Mm01276696_m1) TaqMan Assay | Thermo Fisher Scientific | 4331182 |
| Clca1 (Mm01320697_m1) TaqMan Assay | Thermo Fisher Scientific | 4331182 |
| Alpi (Mm01285814_g1) TaqMan Assay | Thermo Fisher Scientific | 4331182 |

| | | |
|---|---|---|
| Fabp1 (Mm00444340_m1) TaqMan Assay | Thermo Fisher Scientific | 4331182 |
| Chga (Mm00514341_m1) TaqMan Assay | Thermo Fisher Scientific | 4331182 |
| Cav1 (Mm00483057_m1) TaqMan Assay | Thermo Fisher Scientific | 4331182 |
| Cav2 (Mm01129337_g1) TaqMan Assay | Thermo Fisher Scientific | 4331182 |
| Foxc1 (Mm01962704_s1) TaqMan Assay | Thermo Fisher Scientific | 4331182 |
| Gapdh (Mm99999915_g1) TaqMan Assay | Thermo Fisher Scientific | 4448486 |
| **Experimental models: Cell lines** | | |
| HEK293 R spondin-1 | Sigma-Aldrich | SCC111 |
| *Rosa26*-CreER; *Eif4g2*$^{+/+}$, *Eif4g*$^{flox/+}$, and *Eif4g2*$^{flox/flox}$ mouse intestinal organoids | This paper | N/A |
| **Deposited data** | | |
| Raw and analyzed RNA-seq data | This paper | GSE294512 |
| Raw and analyzed Ribo-seq and paired RNA-seq data | This paper | GSE294619 |
| Raw and analyzed data (mass spectrometry) | This paper | jPOSTrepro (https://repository.jpostdb.org/), JPST003517 (PXD059198) |
| **Software and Algorithms** | | |
| Prism 10 | GraphPad | https://www.graphpad.com/scientific-software/prism/ |
| Compass for Simple Western software | Protein Simple | https://www.bio-techne.com/resources/instrument-software-download-center |
| EnrichR | Chen et al[67] | https://maayanlab.cloud/Enrichr/ |
| clusterProfiler (v4.12.6) | Xu et al[68] | https://bioconductor.org/packages/clusterProfiler/ |
| Hisat2 (v2.0.5) | Mortazavi et al[69] | https://github.com/DaehwanKimLab/hisat2 |
| featureCounts (v1.5.0-p3) | Liao et al[70] | https://subread.sourceforge.net/ |
| DeSeq2 (v1.20.0) | Love et al[71] | https://doi.org/10.1186/s13059−014−0550−8 |
| Fastp | Chen et al[72] | https://github.com/OpenGene/fastp |
| Cutadapt 3.7 | Martin et al[73] | https://cutadapt.readthedocs.io/ |
| STAR v2.7.0a | Dobin et al[74] | https://github.com/alexdobin/STAR |
| UMItools v1.1.2 | Smith et al[75] | https://github.com/CGATOxford/UMI-tools |

| | | |
|---|---|---|
| diaPASEF | Meier et al[76] | TimsTOF Pro (Bruker) |
| DIA-NN (v1.8.2 beta27) | Demichev et al[77] | https://github.com/vdemichev/DiaNN |
